# Organelle distribution and protein translation defects in radial glia cells with mutations in Eml1

**DOI:** 10.64898/2025.12.24.696387

**Authors:** V. Viola, K. Chinnappa, R. Thompson, C. Cifuentes Diaz, S. Chakraborty, R. Belvindrah, M. Savariradjane, C. Janke, M.M. Magiera, D. Rinaldi, R. Shayan, V. Soldan, C. Plisson-Chastang, S. Lebaron, M. Cohen-Salmon, F. Francis

## Abstract

Cerebral cortical development is a finely regulated process, depending on key neuronal progenitors, giving rise to post-mitotic neurons that migrate to superficial regions in the developing brain. Disruptions can give rise to severe cortical malformations that are associated with intellectual disability and intractable epilepsy. Mutations in *EML1/Eml1*, coding for a microtubule (MT)-associated protein, lead to subcortical heterotopia (SH) in both humans and mice. In developing mutant mouse brains, ectopic apical radial glia (aRG) progenitors were found outside the normal proliferative ventricular zone (VZ). The normal role of Eml1 in these cells remains to be precisely determined. Using proteomics and imaging approaches, we compared brain tissue from control and mutant mice and identified microtubule and organelle dysregulation. Eml1 influences the trafficking of organelles to different compartments of aRG, where they may be involved in local functions. Thus, we find that mitochondrial and polysome distribution are altered in mutant conditions. Indeed, Eml1 interacts with translational machinery, and an overall downregulation of translation is observed when analysing the translatome of Eml1 mutant aRG. Paradoxically, an upregulation of translation-related mRNAs is also observed. These new data uncover an unprecedented function of Eml1, as well as associating novel subcellular patho-mechanisms with severe heterotopia formation.

## Introduction

Corticogenesis refers to the processes of proliferation, migration, differentiation and synaptogenesis by which the six-layered cerebral cortex is formed in mammals. In mouse, this process starts at E9-10 after telencephalon specification, corresponding to GW6 in humans. At the onset of neurogenesis at E10 (GW6), neuroepithelial cells (NEC) give rise to more fate-restricted progenitors termed apical radial glial cells (aRG). These exhibit both neuroepithelial and astroglial properties. Their cell bodies are restricted to the ventricular zone (VZ), the most apical cell layer that faces the lumen of the cerebral ventricles. aRG have short apical processes, allowing their attachment at the ventricular surface, and a long basal process, reaching the pial surface and acting as a scaffold for migrating neurons (Rakic et al. 1972, Meyerink et al. 2020). These cells can self-renew by symmetric division or give rise to basal or intermediate progenitors (IP), as well as neurons (direct neurogenesis) by asymmetric division (Götz and Huttner 2005).

aRG are highly polarized. Their apical processes terminate with endfeet, helping to form the ventricular boundary. These exhibit a primary cilium (PC) in interphase, protruding into the cerebrospinal fluid and acting as a signalling hub. Longer basal processes span the thickness of the cortex and provide adhesive contacts with migrating neurons (Nadarajah and Parnavelas, 2002). They also terminate with endfeet, contacting the basement membrane at the pial surface. aRG undergo interkinetic nuclear migration during their cell cycle: they are in S-phase when their nuclei are on the basal side of the VZ and in mitosis when they are at the apical side bordering the ventricle (Sauer and Walker, 1959). Establishment and maintenance of aRG cell structure and polarity is crucial for their correct division, organized neuronal migration, and ultimately for proper cortex development. Indeed, several mutations have been shown to impact aRG polarity and function and lead to cortical malformations such as microcephaly and heterotopia (Bizzotto and Francis 2015, Viola et al. 2024).

Mutations in microtubule (MT)-associated proteins (MAPs) can lead to cortical malformations (Romero et al. 2018). Echinoderm Microtubule-associated protein Like 1 (EML1, or EMAPL1) mutations are associated with atypical ribbon-like subcortical heterotopia (SH) (Kielar et al. 2014, Oegema et al. 2019). Patients with compound heterozygous or homozygous *EML1* mutations present ectopic neurons in the white matter, below the cortex, and this is associated with intractable epilepsy, intellectual disability and developmental delay (Kielar et al. 2014, Shaheen et al. 2017, Oegema et al. 2019, Markus et al. 2021). The N terminal domain of EML1, an EMAP family member, binds to MTs, aided by its C-terminal domain, organized in two tandem β-propellers, acting as a protein interaction scaffold (Richards et al. 2014, 2015).

The effects of mutations in *Eml1* were first studied in *Heterotopic Cortex* (*HeCo*) mice, exhibiting a spontaneously arisen retrotransposon insertion in Eml1 intron 22, and leading to the absence of the full-length protein (Kielar et al. 2014). More recently, an Emx1 forebrain-specific conditional knockout (cKO) was created (Zaidi et al. 2024, Vermoyal et al. 2024). This model strongly resembles the *HeCo* model, displaying bilateral SH. Both models consistently show an abnormal delamination of a proportion of aRG, which are therefore found outside the VZ, in the intermediate zone and cortical plate, and proliferate ectopically (Kielar et al. 2014, Uzquiano et al. 2019, Zaidi et al. 2024). Abnormal neuronal migration is likely to be a secondary consequence of altered aRG position. Subcellular defects in aRG were described, affecting the centrosome and MT nucleation (Zaidi et al. 2024). The PC at the apical extremity of aRG does not form correctly, and defects in the Golgi apparatus and post-Golgi trafficking were also described (Uzquiano et al. 2019, Zaidi et al. 2024).

In this study, we revealed novel and unexpected cellular and subcellular defects in Eml1 cKO aRG. Notably, we observed organelle dysregulation using multi-omics and imaging approaches. Mitochondrial and polysome density and distribution are altered in mutant conditions, impacting apical and basal endfeet. Eml1 influences the trafficking of mitochondria to different compartments of aRG, where they may be involved in local functions. We also show that Eml1 interacts with translational machinery components. Furthermore, analysis of the translatome in WT and Eml1 cKO aRG shows overall downregulation of translation in mutant cells. These new data uncover an unexpected role of the MT-binding protein Eml1, as well as associating novel subcellular patho-mechanisms with severe heterotopia formation.

## Results

### Mitochondria transport is altered in Eml1 cKO aRG in culture

Highly polarized cell types such as neurons depend on the MT cytoskeleton for trafficking and distribution of organelles to distal regions of the cell (Maday et al. 2014). Eml1 binds to MTs, and its mutation has already been shown to impact MT polymerization and post-Golgi trafficking in vitro (Bizzotto et al. 2017, Uzquiano et al. 2019, Zaidi et al. 2024, Yigit et al. 2025). Analysing previously generated interactome and proteome data (Bizzotto et al. 2017, Uzquiano et al. 2019, Zaidi et al. 2024, Yigit et al. 2025), we found that proteins related to transport and trafficking are downregulated in Eml1 cKO brains (Table 1). Mitochondrial transport has been widely studied and is known to rely on the MT cytoskeleton, as well as kinesin and dynein motors (Melkov and Abdu 2018, Schwarz 2013). Rare studies observing mitochondria and their movement in aRG have been published, when studying metabolism (Rash et al 2018). We decided therefore to further assess trafficking in the Eml1 cKO condition, focusing on mitochondria, which are likely to be transported in polarised aRG and play local functions.

**Table 1.**
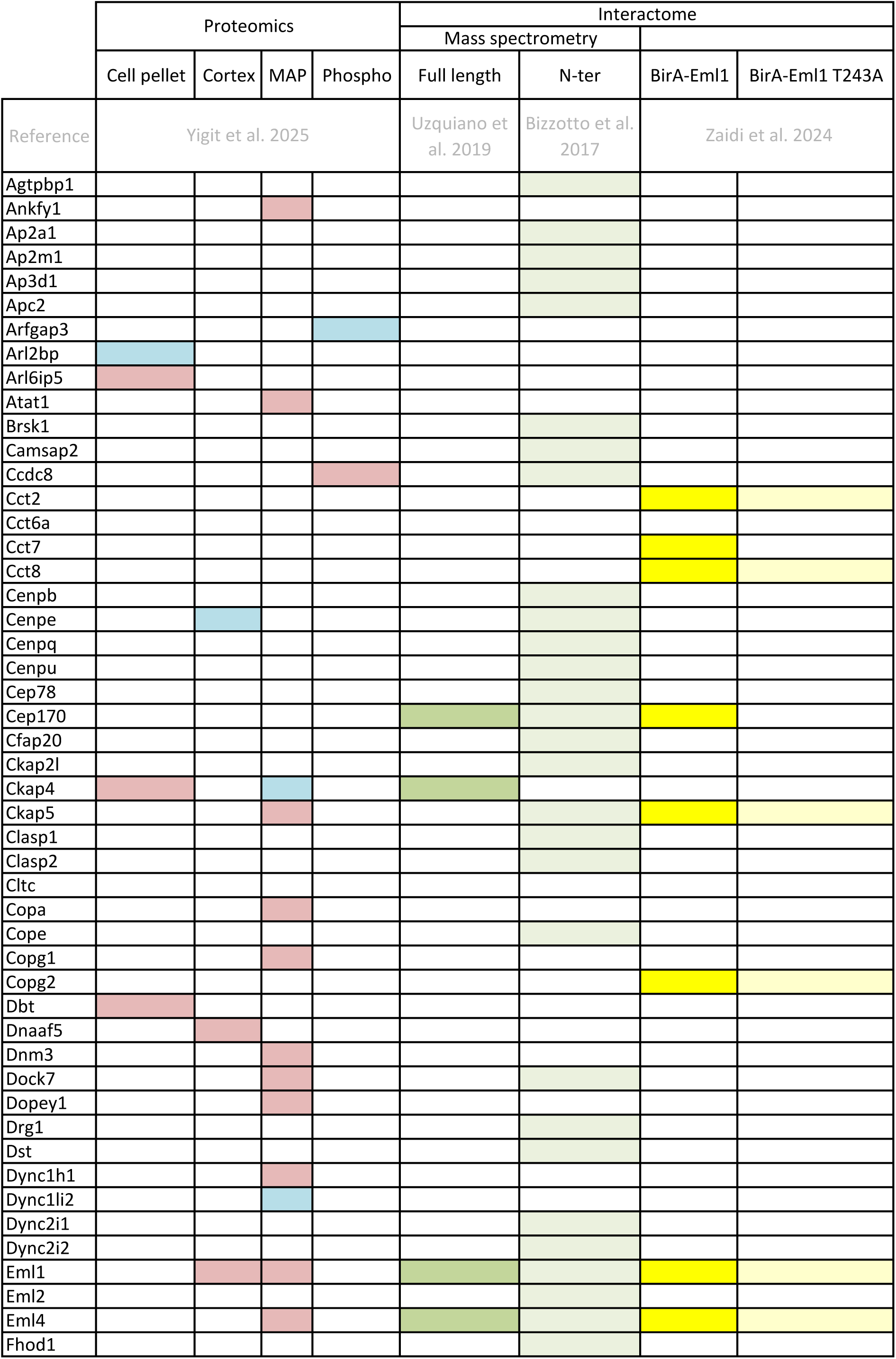
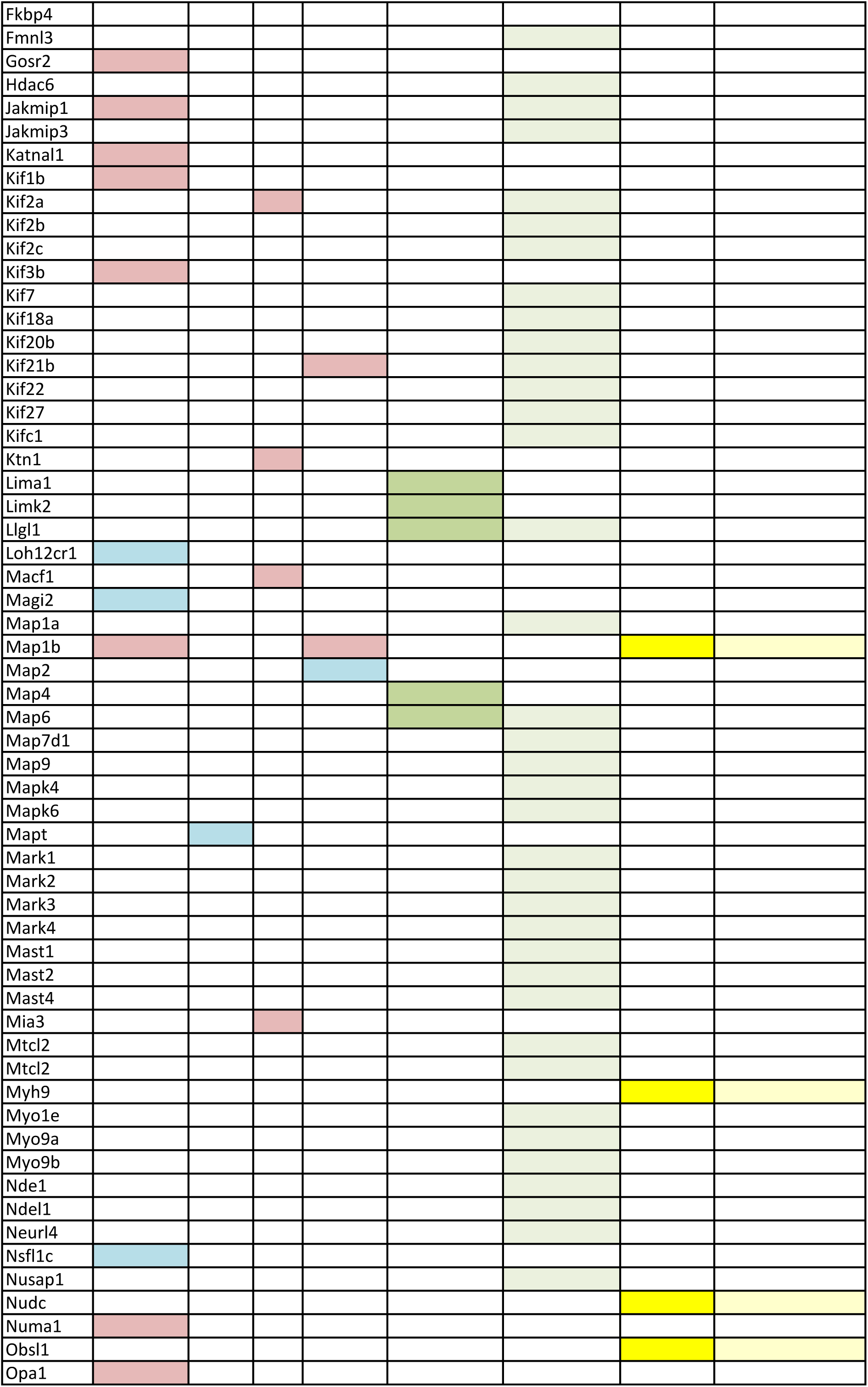
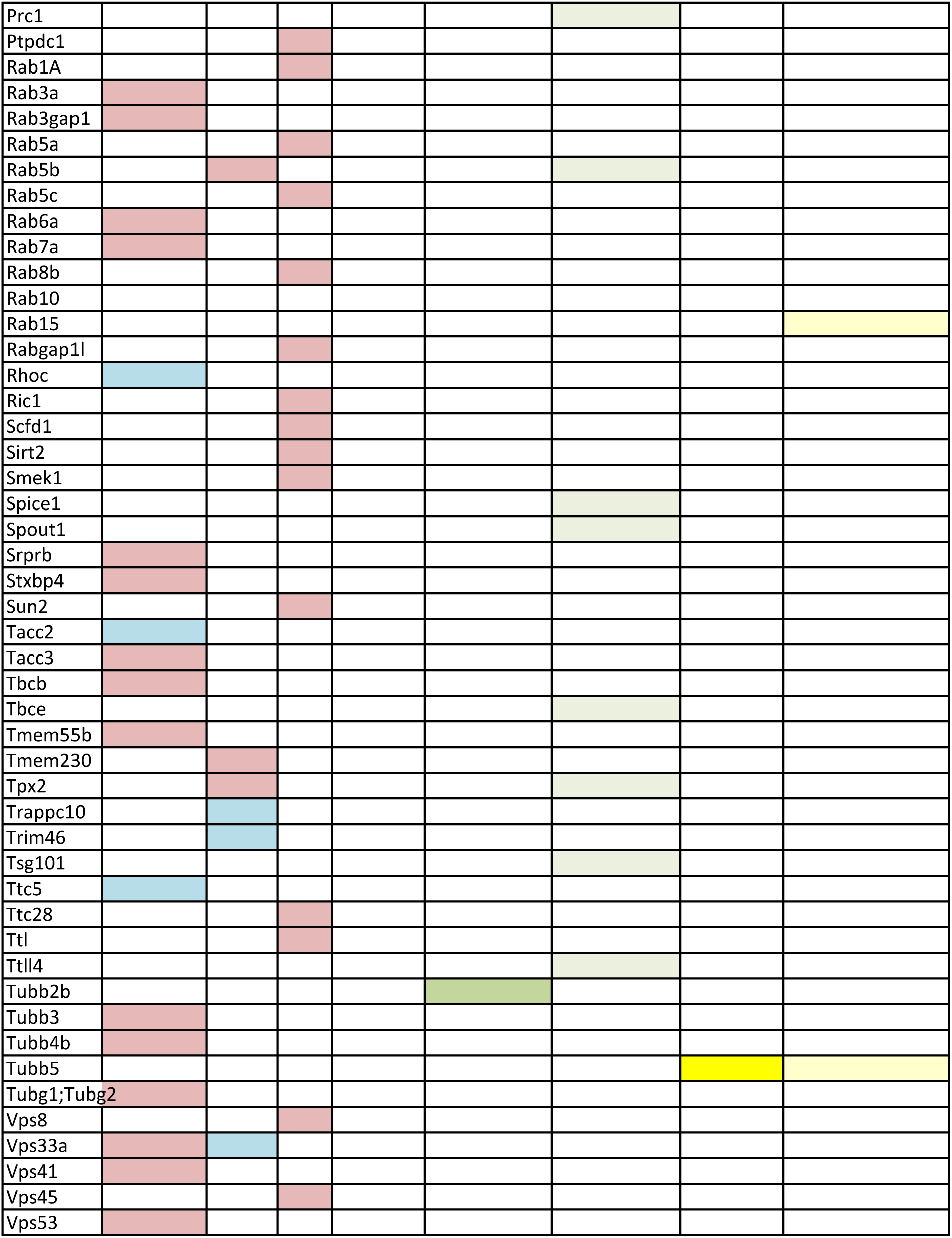
Proteins related to MT cytoskeleton and transport in proteomic and interactome screens. . Blue: upregulated, red: downregulated.

Primary cultures of aRG-like progenitor cells from WT and Eml1 cKO embryonic cortices were generated (Sun et al. 2011). Pax6+ aRG in culture have processes that extend from the cell soma, although they lack apico/basal polarity as observed in vivo. Centrosomes are observed close to the nucleus in the cell soma (Supplementary Figure 1A). Cells were exposed to MitoTracker and live imaging was performed. One process per cell was selected and kymographs obtained to measure mitochondria mobility parameters (Figure 1A, B). Eml1 cKO mitochondria showed increased speed compared to WT, in particular in the inward direction towards the soma. Pausing time was reduced, contributing to the increased speed of Eml1 cKO mitochondria compared to WT (Figure 1C). Distance and persistence parameters were not significantly altered (Figure 1D, Supplementary Figure 1B). Thus, mitochondria appear to move faster towards the somal MT organizing centre (in vivo this is found in aRG apical endfeet during interphase, Wimmer and Baffet 2023).

**Figure 1.**
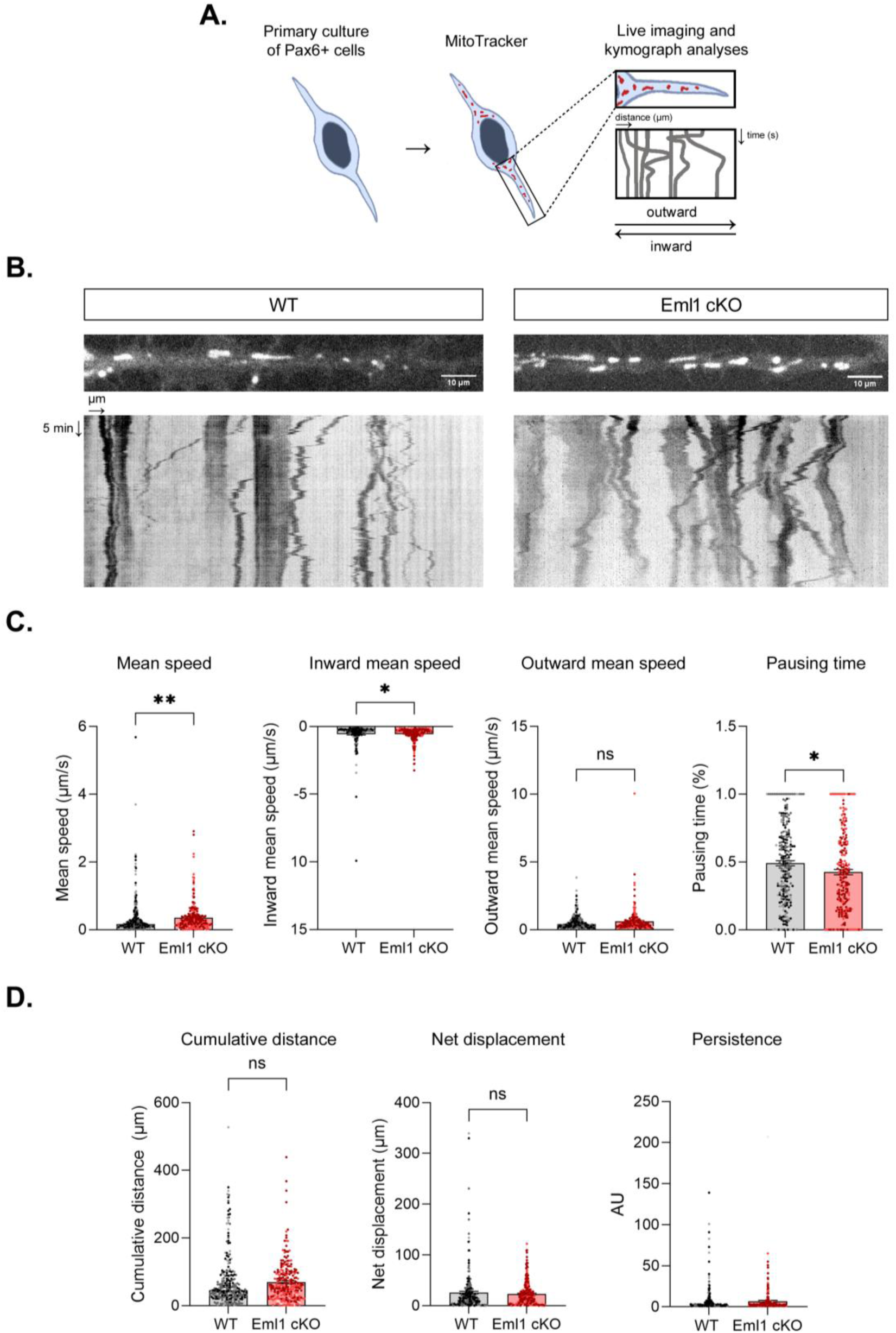
Mitochondria transport is affected in Eml1 cKO aRG. **A.** Workflow of MitoTracker treatment of primary cultures of Pax6+ cells. Cells are exposed for 30 minutes to stain mitochondria, and imaged directly after with a spinning disk microscope. A single process per cell was analysed with KymoToolBox to obtain a kymograph, showing X axis = distance and Y axis = time. Cell soma to process is defined as outward, process to soma as inward transport. **B.** Representative snapshot of WT and Eml1 cKO Pax6+ cell process stained with MitoTracker, and their respective kymographs. **C.** Quantification of mean speed, inward mean speed, outward mean speed and pausing time of WT and Eml1 cKO mitochondria. Values are represented as mean ± SEM. N = 241 (WT) and 246 (Eml1 cKO) mitochondria, 26 cells analysed per genotype from 3 different cultures, indicated by dots of different colours in the graphs. Mann-Whitney test, p-value ** ≤0.01, * ≤0.05. **D.** Quantification of cumulative distance, net displacement and persistence of WT and Eml1 cKO mitochondria. Values are represented as mean ± SEM. N = 241 (WT) and 246 (Eml1 cKO) mitochondria, 26 cells analysed per genotype from 3 different cultures, indicated by dots of different colours in the graphs. Scale bar, equivalent for WT and Eml1 cKO: 10 µm.

Tubulin post-translational modifications (PTMs) are important regulators of MT functions, influencing processes such as intracellular transport and interactions with MAPs and motors (Janke and Magiera, 2020). Polyglutamylation, one of the tubulin PTMs prevalent on neuronal MTs, has been shown to regulate mitochondrial transport (Magiera et al. 2018, Gilmore-Hall et al. 2019, Bodakuntla et al. 2020, Bodakuntla et al. 2021) and MAP interaction with MTs (Genova et al. 2023). To evaluate if the transport phenotype could be linked to changes of tubulin PTM levels in Eml1 cKO mouse brain, we used electrochemiluminescence (ECL)-based immunoassays, which allow for a quantitative analysis of tubulin PTMs. We analysed protein lysates of WT and Eml1 cKO E12.5 cortices (one day before aRG delamination has been observed in this model, Zaidi et al. 2024) and extracts of cultured Pax6+ progenitor cells, examining several key tubulin PTMs: tyrosination, detyrosination, polyglutamylation and acetylation. Although a mild reduction in total β-tubulin levels and a trend toward increased PTM levels in Eml1 cKO samples was observed, none of these differences were significant (Supplementary Figure 1C, D).

These combined data point to changes in mitochondria trafficking in aRG, apparently unrelated to the tubulin PTM status.

### Mitochondria density is increased in Eml1 cKO apical, and decreased in basal endfeet, at E12.5 and E15.5

We further questioned organelle position in mutant aRG in brain sections. Mitochondria are known to be broadly distributed in aRG, including endfeet (Rash et al. 2018). Performing electron microscopy (EM), images were analysed from E12.5 and E15.5 embryonic brains. Apical endfeet were identified by their proximity to the ventricles, and basal endfeet were identified by their proximity to the cortical pial surface, where the basal lamina and the meninges were clearly visible (Figure 2A). WT aRG presented mitochondria in both apical and basal endfeet, but these were enriched in the latter at E12.5 (Figure 2B, 2D), with relatively fewer observed at E15.5 (Figure 2C, 2E). In Eml1 cKO aRG mitochondria density was significantly increased in apical endfeet at both stages. Significantly decreased density was observed in E12.5 basal endfeet in Eml1 cKO aRG, with a similar tendency for reduction observed at E15.5 (Figure 2D, 2E). This leads to an inversed distribution of mitochondria in Eml1 cKO aRG extremities at E12.5 (higher in apical and lower in basal), with similar trends at E15.5. For apical endfeet when comparing the two stages, mitochondria densities appear similar in WT and are similarly increased in the cKO at both stages, whereas in basal endfeet a notably higher density is observed at E12.5 compared to E15.5 in WT, with no differences in the cKO across the stages (Supplementary Figure 2A). Mitochondrial element morphology such as area, perimeter, circularity, length and width were overall not different at E12.5 and E15.5 (Supplementary Figure 2B, C), but some tendencies for reduced area, perimeter and width were observed in WT mitochondria elements in basal endfeet compared to apical endfeet, reaching significance at E15.5 (Supplementary Figure 2C).

**Figure 2.**
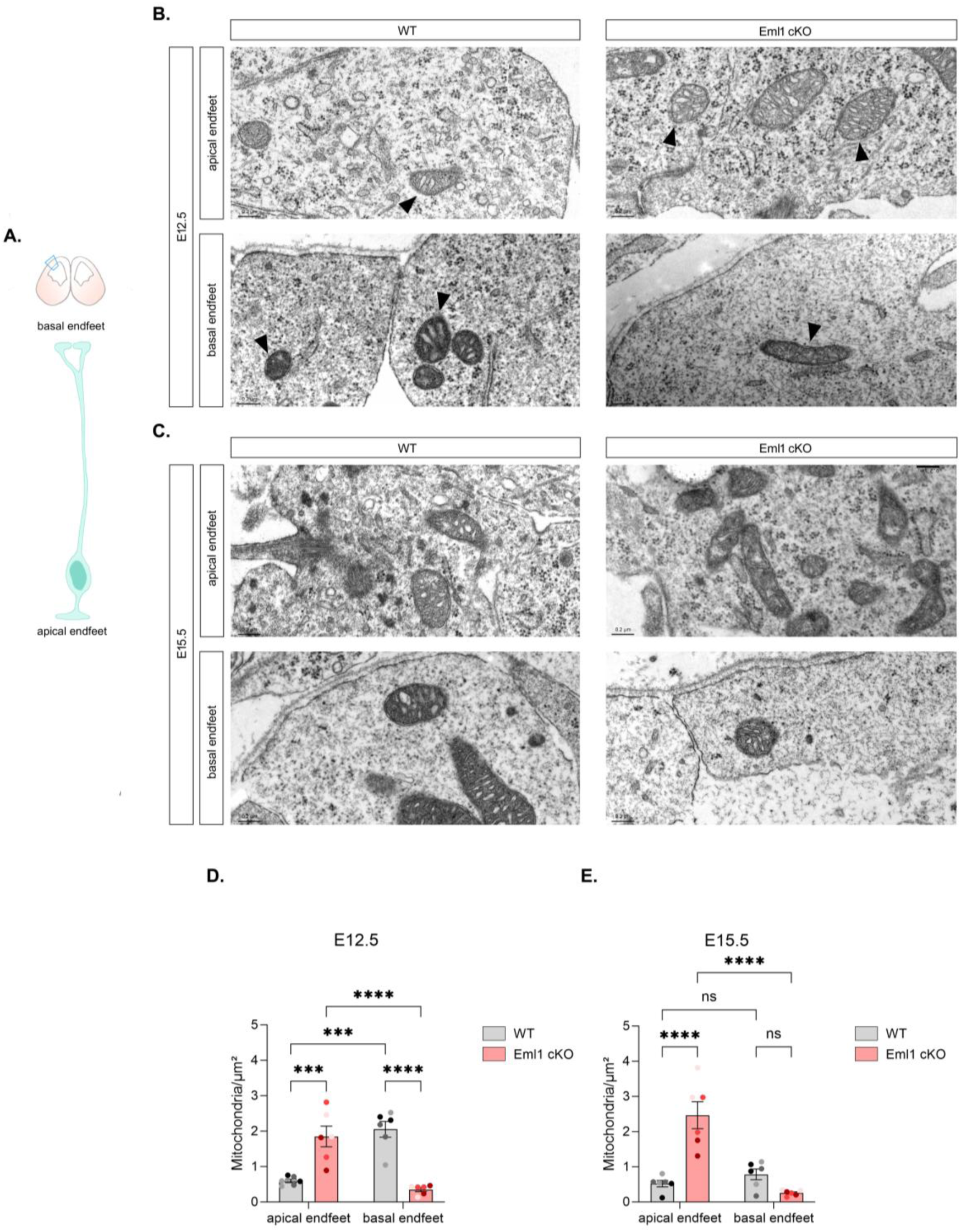
Mitochondrial element density is increased in apical endfeet, decreased in basal endfeet of Eml1 cKO radial glia cells. **A.** Schema depicting a coronal section of mouse embryonic cortex and an aRG, indicating apical and basal endfeet. **B, C.** Electron microscopy images of E12.5 (**B**) and E15.5 (**C**) aRG apical and basal endfeet, where mitochondrial elements can be observed (black arrowheads). **D, E.** Quantification of mitochondrial element density in E12.5 (**D**) and E15.5 (**E**) apical and basal endfeet. Values are represented as mean ± SEM. N = 6 embryos per genotype from 3 different litters, indicated by dots of different colours in the graphs. Two-way ANOVA, p-value **** ≤0.0001, *** ≤0.001. Scale bar, equivalent for WT and Eml1 cKO: 0.2 µm.

EM analyses thus reveal endfeet specific mitochondria features in polarised aRG at different stages, and show in the mutant, altered distribution, with mitochondria increased in apical and decreased in basal endfeet in Eml1 cKO aRG compared to WT.

### Eml1 interacts with ribosomes and translational machinery

The Eml1 ancestral protein EMAL was first described in sea urchin eggs (Suprenant et al. 1993). As MT preparations decorated with ribosomes changed upon EMAL protein removal (Suprenant et al. 1993), this protein was evoked as a potential MT-ribosome adaptor.

Our Eml1 interactome experiments and gene ontology (GO) analyses also indicated that the biological processes of translation, peptide biosynthetic process and peptide metabolic process were enriched when assessing EML1 proximal interactors in transfected neural cells (Zaidi et al. 2024). Further analysing these data, we identified 16 out of 50 (N-terminal BioID tag, N term) proximal interactors (and 17 out of 58, C-terminal BioID tag, C term interactors, data not shown) corresponding to ribosomal proteins or proteins involved in ribosome biogenesis and function (Table 2, Figure 3A). Importantly, more than half of these proteins (9) lost their interaction when EML1 carried the patient mutation T243A (and 11 for C term interactors, data not shown) (Figure 3A). Ribosomal proteins of both large (Rpl) and small (Rps) subunits were identified, such as Rps3, Rps27 and Rpl17, as well as translation initiation factors (Eif2s3x, Eif4b, Eif4g1).

**Figure 3.**
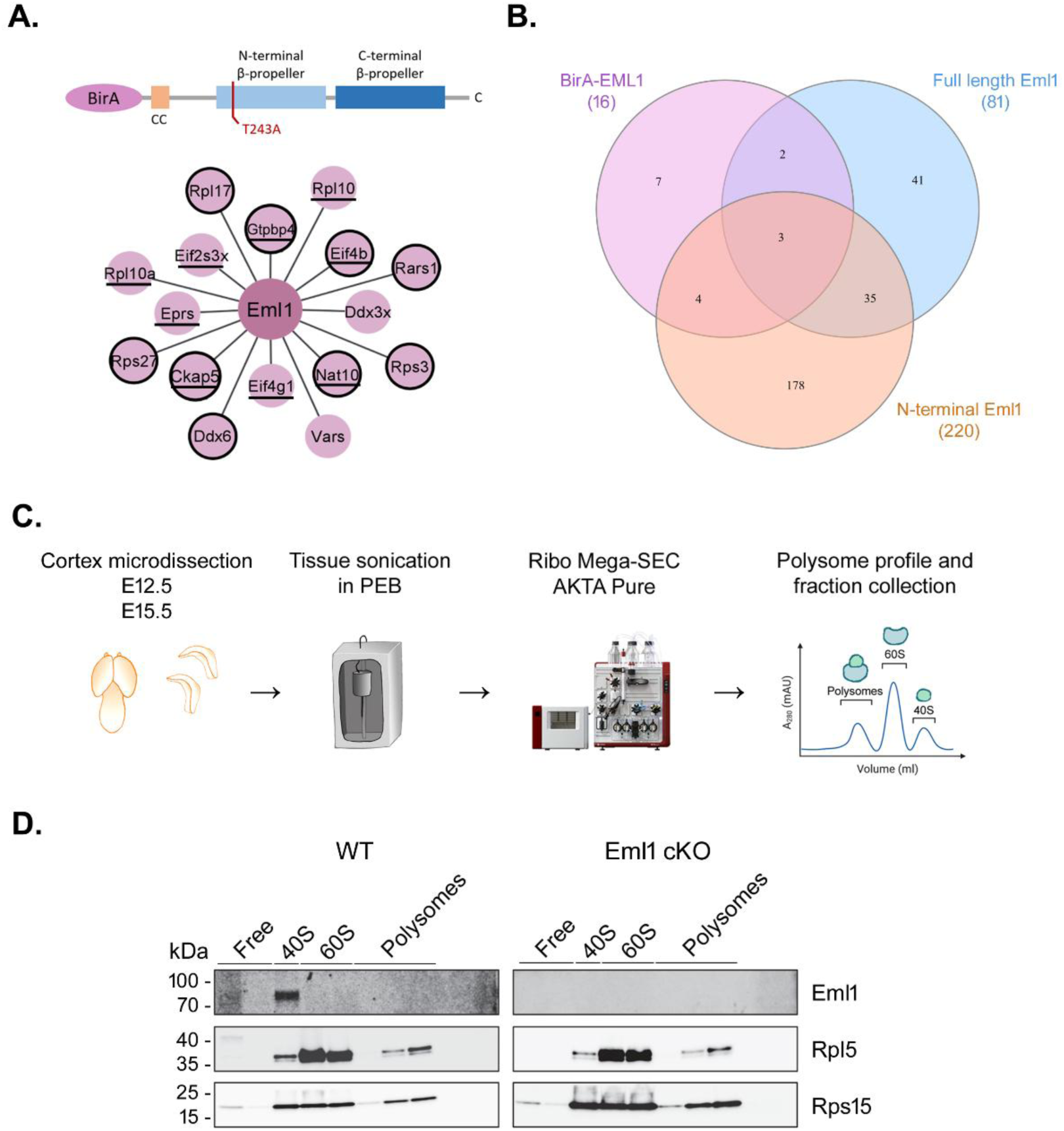
Eml1 interacts with translational machinery components. **A.** Ribosomal proteins, ribosome biogenesis and translation related proteins were identified in EML1-BirA (data not shown) and BirA-EML1 (right) BioID interactor screens. Circled proteins lose interaction with EML1 carrying the patient mutation T243A. **B.** Venn diagram showing intersections between different datasets of Eml1 interactome experiments (including previous N terminal and full-length screens, Bizzotto et al. 2017, Uzquiano et al. 2019) for ribosomal proteins, ribosome biogenesis and translation-related proteins. **C.** Workflow of Ribo Mega-SEC experiments for embryonic cortices. Brains were microdissected to isolate the cortex at different embryonic stages. The tissue was lysed with a sonicator in a polysome extraction buffer (PEB). The lysate was then run on an AKTA Pure machine, and a polysome profile and collectable fractions for the different subunits were obtained. **D.** Western blot of ribosomal fractions from E15.5 cortices obtained after Ribo Mega-SEC. Note that Eml1 is found in the 40S fraction. For comparison, Rpl5 and Rps15 proteins were tested. N = 3 experiments from 3 different litters, 3 embryos were pooled per genotype.

**Table 2.**
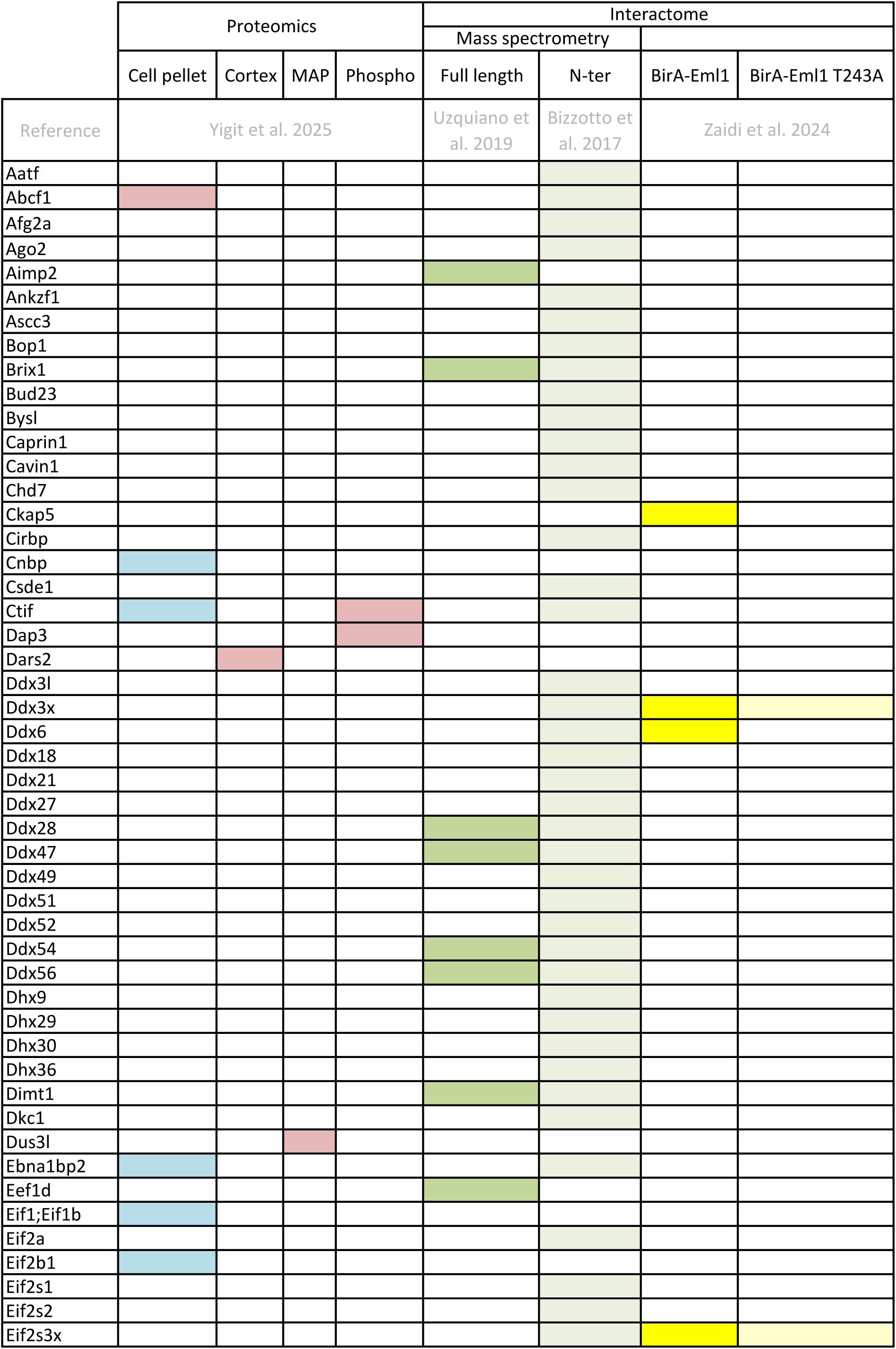
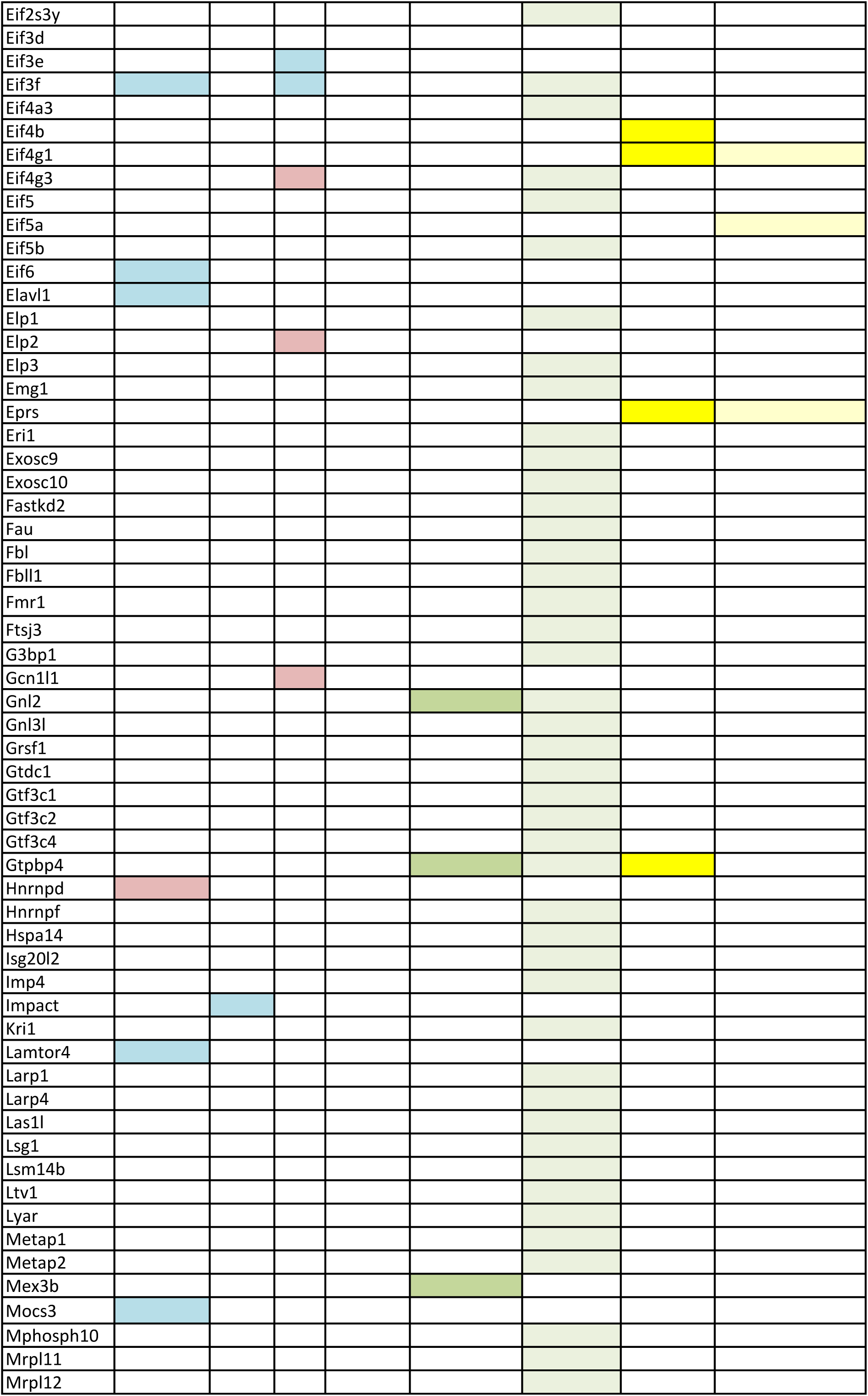
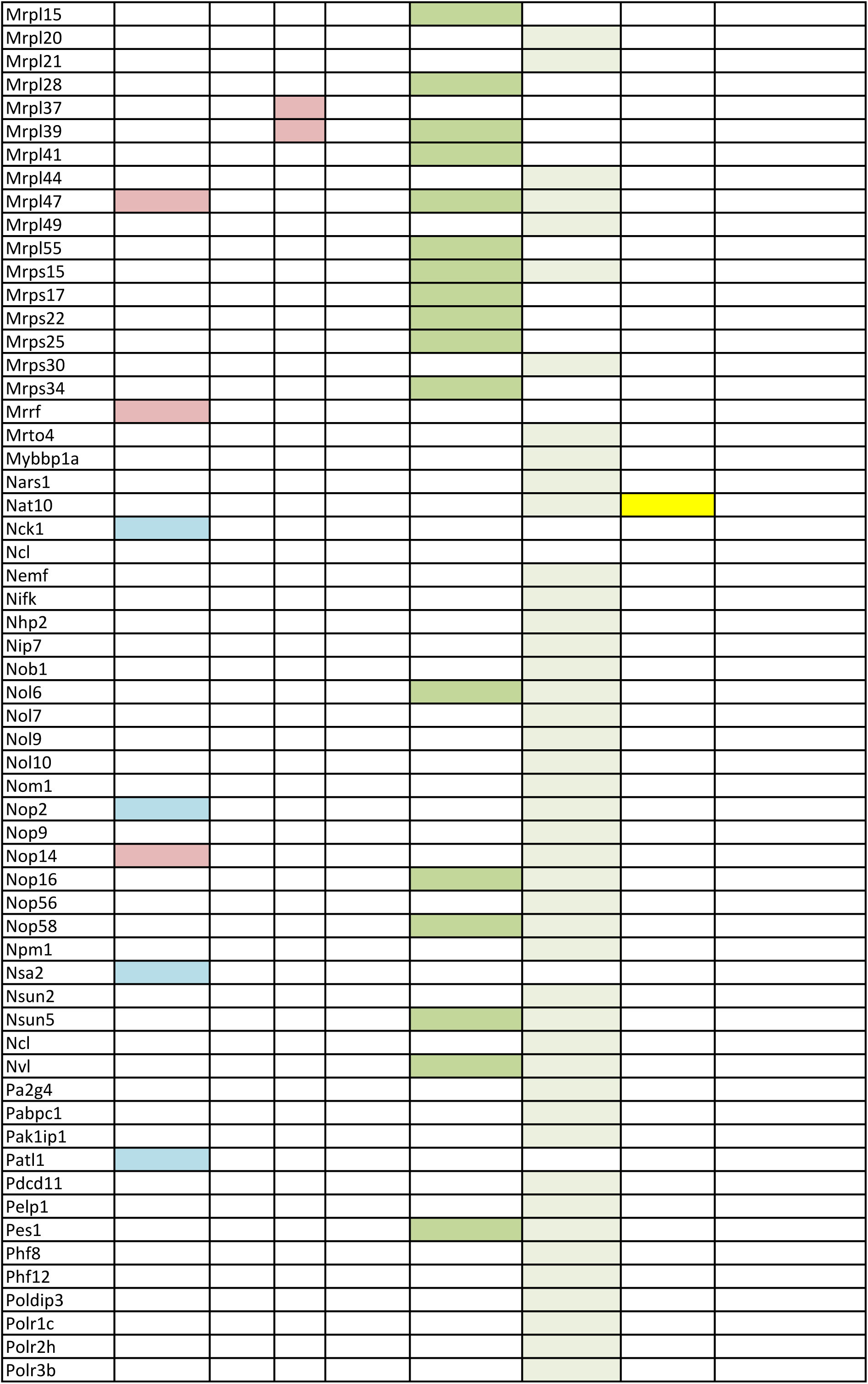
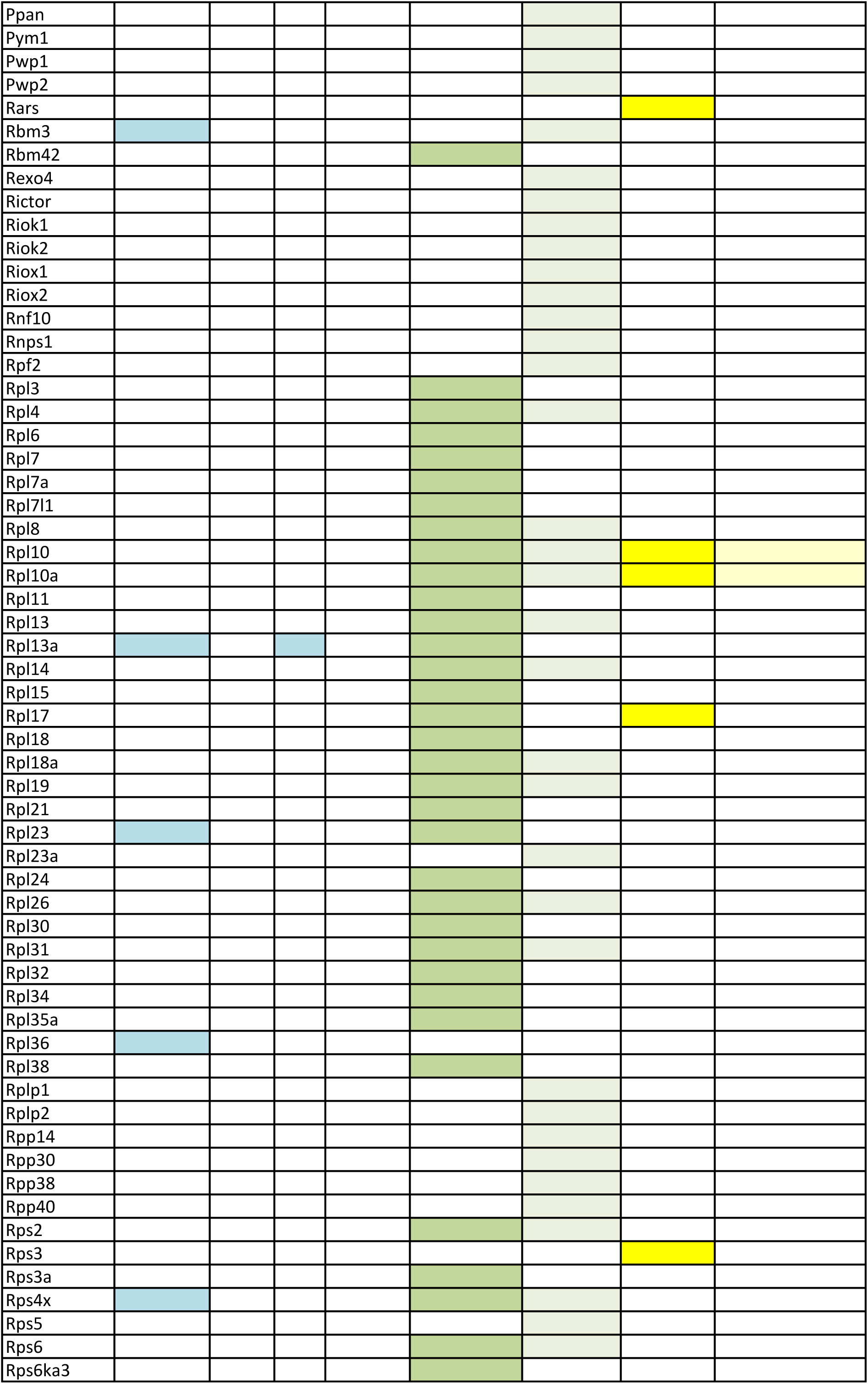
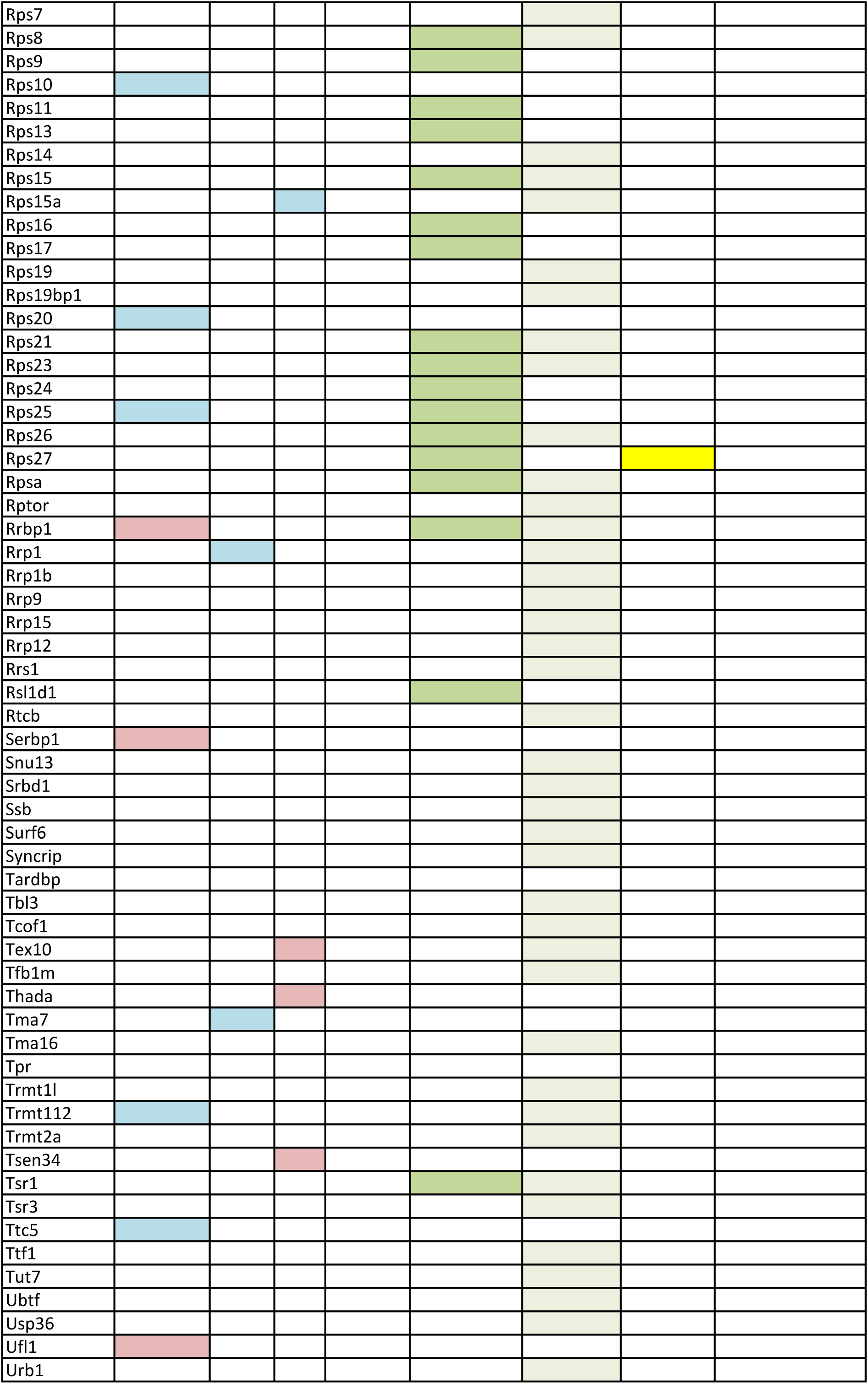
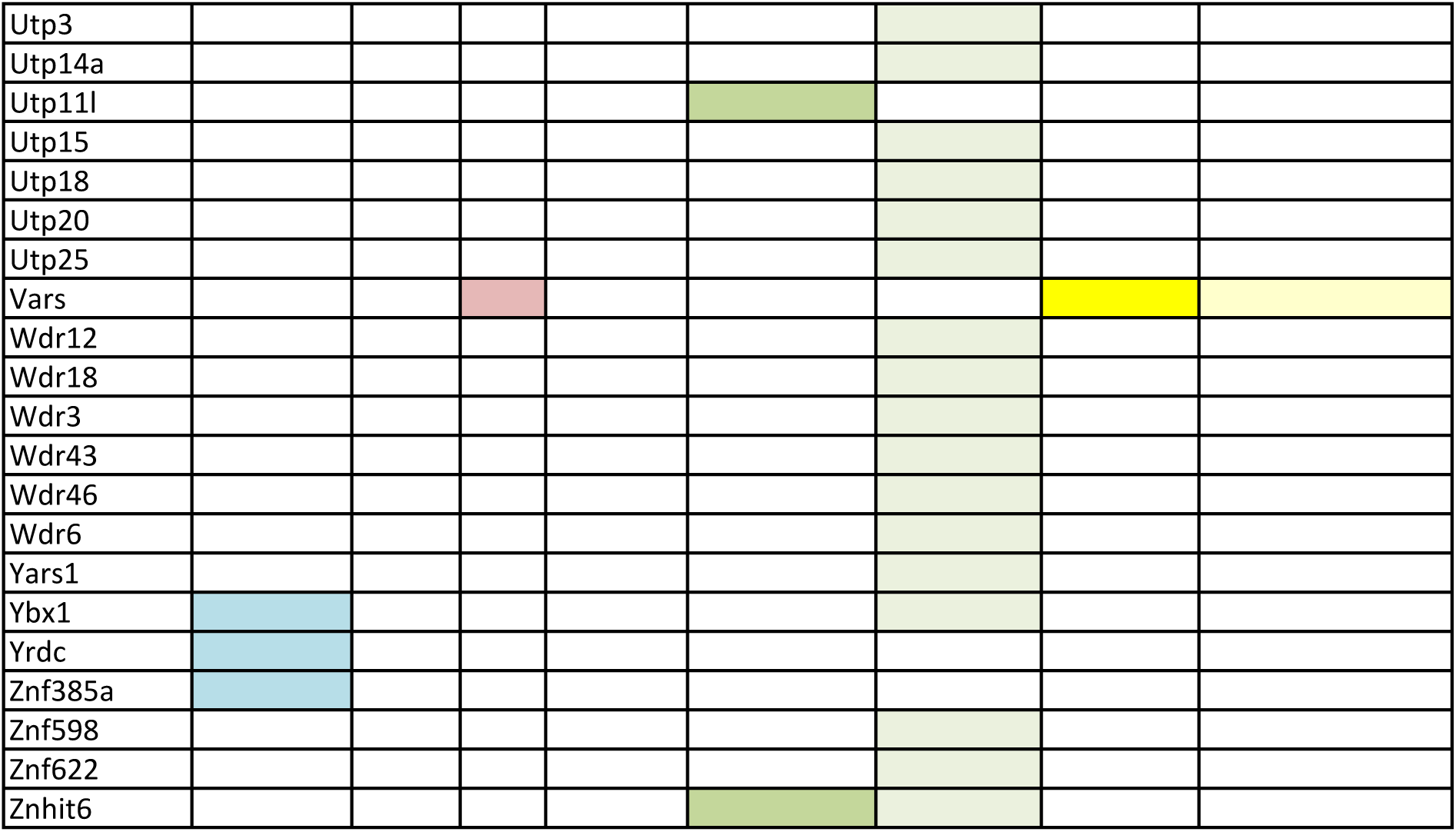
Proteins related to ribosome and translation in proteomic and interactome screens. Blue: upregulated, red: downregulated.

These data overlap with previously performed screens for Eml1 interactors identified in mouse embryonic cortex extracts (Bizzotto et al. 2017, Uzquiano et al. 2019) (Figure 3B, Table 2). This was the case for ribosome-related interactors, as well as ribosomal proteins Rpl10, Rpl10a, Rpl17, Rps27 and ribosome biogenesis factor Nat10. Taken together, these data suggest a disease-relevant interaction of Eml1 with components of the translation machinery.

To further assess whether Eml1 is associated with ribosomal subunits and/or polysomes, ribosome fractionation was performed using Ribo Mega-SEC approach (Yoshikawa et al. 2018). E12.5 (data not shown) and E15.5 mouse cortices were lysed in a polysome extraction buffer (PEB), prior to running the samples through the AKTA Pure system, which provides a polysome profile and collectable fractions (Figure 3C). Fractions were collected corresponding to 40S and 60S subunits, as well as polysomes, and separated on SDS-PAGE gels. Western blot experiments showed an enrichment of Eml1 signal in 40S, but not in 60S and polysome fractions (Figure 3D). This suggests a preferential interaction of Eml1 with the 40S subunit. Large subunit (Rpl) proteins identified in the interactome may thus associate indirectly within the ribosomal protein complex or correspond to a more transient/labile interaction between Eml1 and large ribosomal subunit.

### Polysome density is increased in Eml1 cKO apical, and decreased in basal, endfeet, at E12.5 and E15.5

The interaction of Eml1 with ribosomal subunits and translation factors may influence their trafficking within the cell. Indeed, they may be moved along MTs in order to exert their functions in distal regions (Pilaz et al. 2016, Chudinova and Nadezhdina 2018). EM was hence used to assess ribosome/polysome distribution in aRG extremities in the developing cortex at E12.5 and E15.5. As for mitochondria, apical endfeet were imaged facing the ventricle and basal endfeet facing the meninges (Figure 4A). Several ribosomes can translate the same mRNA, and thus form a ”beads on a string“ arrangement, known as polysomes (Afonina and Shirokov 2018). Polysomes were clearly visible in both cell compartments (Figure 4A, B). Polysome density was quantified, and we observed that E12.5 WT aRG had a similar polysome density in the apical and basal endfeet. In Eml1 cKO brains, polysome density was increased in apical, and decreased in basal endfeet, creating a difference in density between the two compartments (Figure 4C). In WT aRG at E15.5, contrary to E12.5, the density between the two compartments differed, as lower polysome density was observed in basal compared to apical endfeet in the WT condition (Figure 4D). In Eml1 cKO aRG at E15.5, polysome density was similarly increased in apical (although to a slightly lesser extent compared to E12.5, Supplementary Figure 3A) and decreased in basal endfeet (Figure 4D).

**Figure 4.**
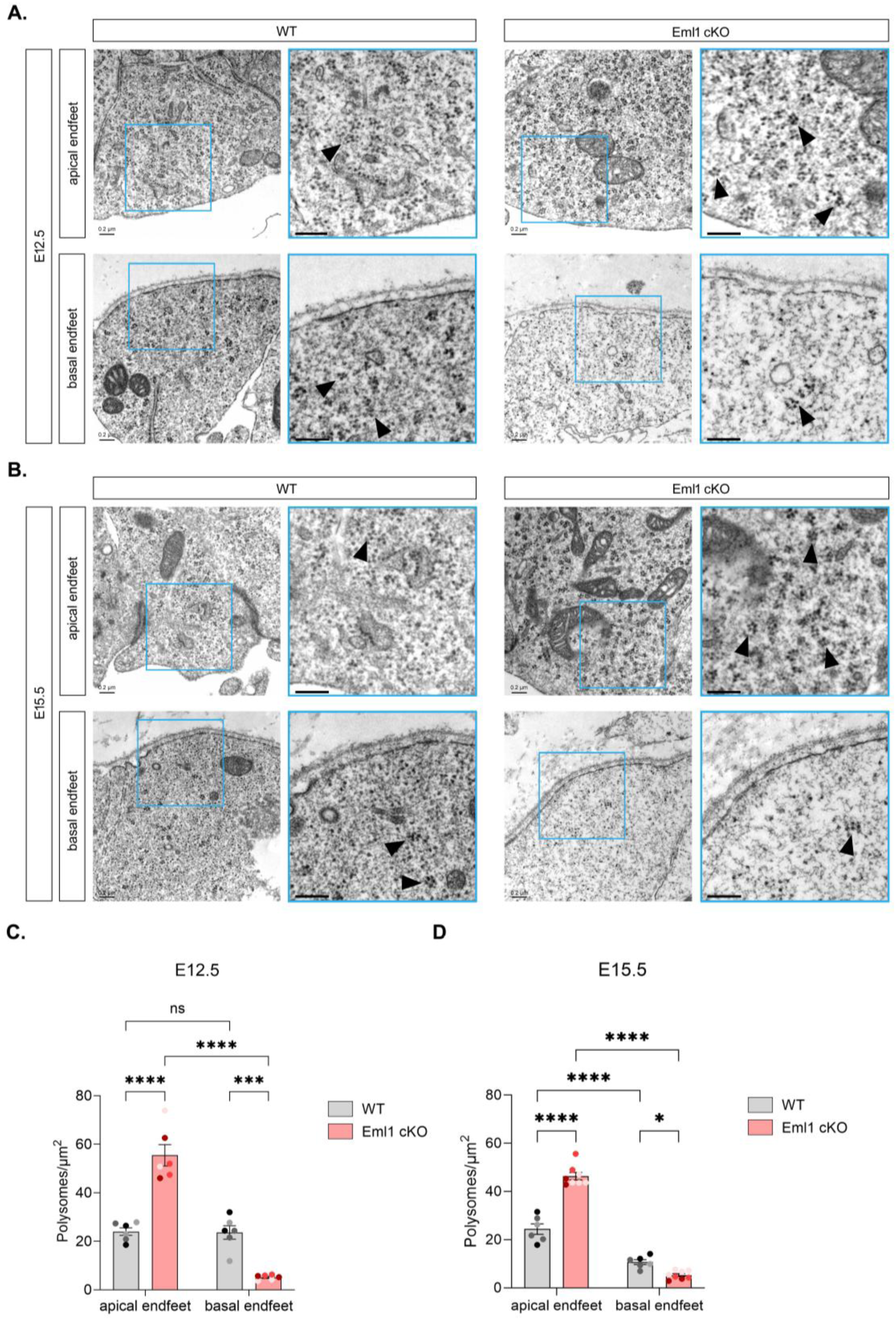
Polysome density is increased in apical and decreased in basal endfeet of Eml1 cKO radial glia cells. A, **B.** Electron microscopy images of E12.5 (**A**) and E15.5 (**B**) aRG apical and basal regions, adjacent to the ventricle and pial surface respectively. Examples of polysomes are indicated by black arrowheads. **C.** Quantification of polysome density in E12.5 apical and basal endfeet. Values are represented as mean ± SEM. N = 6 embryos per genotype from 3 different litters, indicated by dots of different colours in the graphs. Two-way ANOVA, p-value **** ≤0.0001, *** ≤0.001. **D.** Quantification of polysome density in E15.5 apical and basal endfeet. Values are represented as mean ± SEM. N = 6 (WT) and 8 (Eml1 cKO) embryos per genotype from 3 different litters, indicated by dots of different colours in the graphs. Two-way ANOVA, p-value **** ≤0.0001, * <0.05. Scale bar, equivalent for WT, Eml1 cKO and magnifications: 0.2 µm.

To complement these 2D observations, we performed electron tomography on the apical and basal endfeet of both WT and Eml1 mutant HeCo embryonic mouse cortices at E13.5. Tomographic 3D reconstructions were combined with semi-automated segmentations of ribosomes and other cellular components, including mitochondria, MTs, and cytoplasmic and internal membranes (Supplementary Figure 4A). We quantified the volume of segmented ribosomes and polysomes across the tomograms, which revealed a significant increase in apical endfeet of Eml1 mutant compared with WT cortices, and conversely, a decrease in the basal endfeet of Eml1 mutant relative to WT (Supplementary Figure 4B). Notably, the number of ribosomes appears comparable between apical and basal endfeet in mutant mice, whereas in WT mice it is significantly higher in basal than in apical endfeet. Altogether, these results indicate an alteration in polysome density and distribution within aRG compartments during corticogenesis in Eml1-deficient brains.

### aRG endfeet structure are altered, potentially due to abnormal organelle content

aRG endfeet are specialized compartments, contributing to cell polarity, with the apical side facing the ventricle and the basal extremity in contact with the meninges. Each compartment shows a different structure and organelle composition, as well as function (Weimer et al. 2009, Shao et al. 2020, see also Viola et al 2024). We therefore decided to assess if the abnormal distribution of polysomes and mitochondria could have an impact on apical and basal endfeet structure.

Concerning the apical side, adherens junctions (AJ) were first analyzed in EM acquisitions. Shorter AJ were observed at E12.5 in the cKO (Figure 5A, B), which could potentially be associated with AJ weakening. This stage is just prior to excessive delamination (Zaidi et al. 2024). This difference was not detected at E15.5, when aRG delamination is no longer significantly different compared to WT (Figure 5C, D) (Zaidi et al. 2024). Apical endfeet size was also assessed. En face imaging was performed after staining with Phalloidin. This allowed us to identify cell borders at the apical surface and to measure their area (Figure 5E, F). In Eml1 cKO at E13.5, a reduced proportion of apical elements with a diameter smaller than 3 μm, together with an increased proportion of apical elements with a diameter greater than 7 μm, was observed (Figure 5F). Apical element morphology hence appears enlarged in Eml1 cKO conditions.

**Figure 5.**
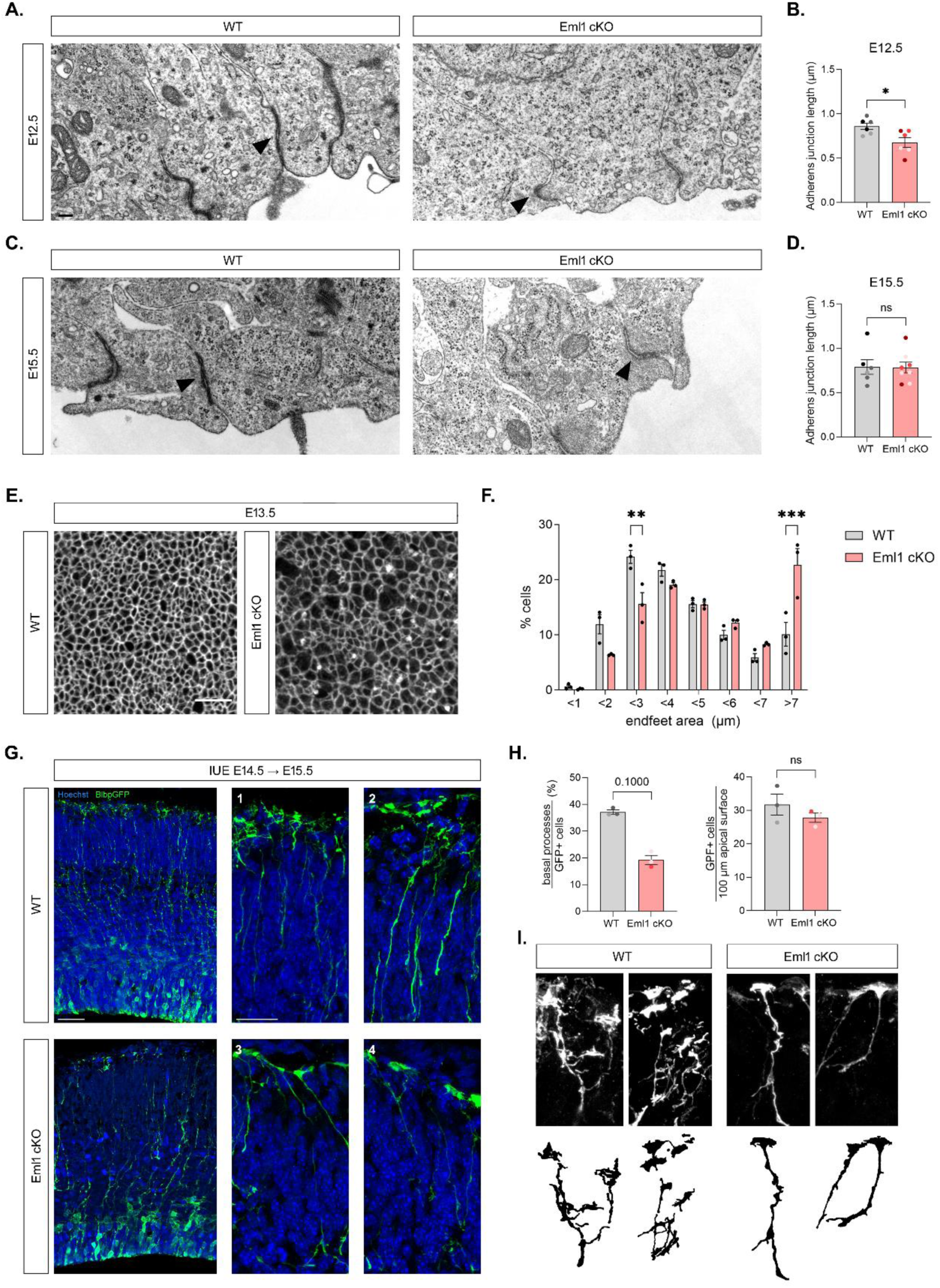
Aspect of aRG extremities. A, **C.** Electron microscopy images of E12.5 aRG apical regions, adjacent to the ventricle, where AJ are visible. **B, D.** Quantification of AJ length. N = 6 embryos per genotype from 3 different litters, indicated by dots of different colours in the graphs. Values are represented as mean ± SEM. Unpaired t-test. Scale bar, equivalent for WT and Eml1 cKO: 0.2 µm. **E.** Apical elements are enlarged at E13.5 in the cKO, as shown by phalloidin staining of boundaries and en face imaging. **F.** Quantification of apical element area. N = 3 embryos per genotype from 3 different litters, indicated by dots of different colours in the graphs. Values are represented as mean ± SEM. Two-way ANOVA, p-value *** ≤0.001, ** ≤0.01. Scale bar, equivalent for WT and Eml1 cKO: 0.2 µm. **G.** Basal processes and endfeet in WT and Eml1 cKO after Blbp-GFP electroporation. Scale bar, equivalent for WT and Eml1 cKO: 40 µm, magnifications: 20 µm. **H**. Quantification of number of processes and GFP+ cells per 100 µm of electroporated area. 3 embryos per genotype from 2 different litters. **I**. Representative images of basal endfeet are schematized for WT and Eml1 cKO.

For basal endfeet morphology, in utero electroporation (IUE) was performed at E14.5 to introduce a brain lipid binding protein (BLBP, also known as Fabp7) reporter (BLBP-GFP) plasmid in aRG in the VZ. One day after IUE, embryos were sacrificed, and confocal microscopy images were acquired to assess GFP+ basal process and endfoot features. In Eml1 cKO, despite similar efficiency of electroporation, less GFP+ basal processes were observed, with less endfeet reaching the pial surface (Figure 5G, H). Also, the complexity of basal endfeet appeared to be reduced, since less ramifications were obvious compared to WT aRG (Figure 5I). Reduced mitochondria and polysomes in the cKO basal endfeet, and presumably in the basal processes as well, might contribute to a reduced attachment and stability of the basal processes (Rash et al. 2018).

Thus, multiple structural apical and basal changes are observed, potentially pointing to altered function.

### Ribosome biogenesis appears increased in Eml1 cKO cortices

As well as endfeet features, we wondered whether overall ribosome/polysome amounts, related to their biogenesis, were also affected, in addition to their altered distribution in the cells. Indeed, Eml1 also associates with factors involved in ribosome synthesis (Nat10, Brix1) (Figure 3A).

To check whether ribosome biogenesis was increased, impacting RNA polymerase I activity and ribosome synthesis, fibrillarin (Fbl) staining was first performed as an indicator of ribosome biogenesis activity in the nucleoli (Baker et al. 2013). We did not observe any differences in Fbl staining between WT and Eml1 cKO, first suggesting that ribosome synthesis is not dysregulated in Eml1 cKO, or that this dysregulation is not strong enough to be detected by this immunostaining procedure (Supplementary Figure 3D, E).

We further assessed the maturation of pre-ribosomal rRNAs (pre-rRNAs) by Northern Blot experiments performed using total RNA extracted from cortices used for Ribo Mega-SEC analysis. The ratios between all pre-rRNA intermediates were quantified relative to WT in order to detect a change in processing (schematized in Figure 6A). At E12.5, the processing appears mostly unaffected except for an accumulation of the primary transcript 47S compared to 32S, which corresponds to the first pre-60S maturation pathway intermediate (Figure 6B, C). We also observed a mild 29S accumulation compared to 34S pre-RNA species. Interestingly, this does not alter mature rRNA species 18S and 28S accumulation (Figure 6B, C). At E15.5, the ratios between the first cleavage products (34S and 32S) and the primary transcript (47S) are significantly reduced (Figure 6D, E). The accumulation of 29S pre-RNA can no longer be observed at this age. The accumulation of 47S compared to the cleavage 2c downstream products 32S and 34S (Figure 6A) observed at both E12.5 and E15.5, can either result from an impaired 2c cleavage, that is only mildly affected and not fully blocked, or by an increase in RNA Polymerase I activity and 47S neo-synthesis.

**Figure 6.**
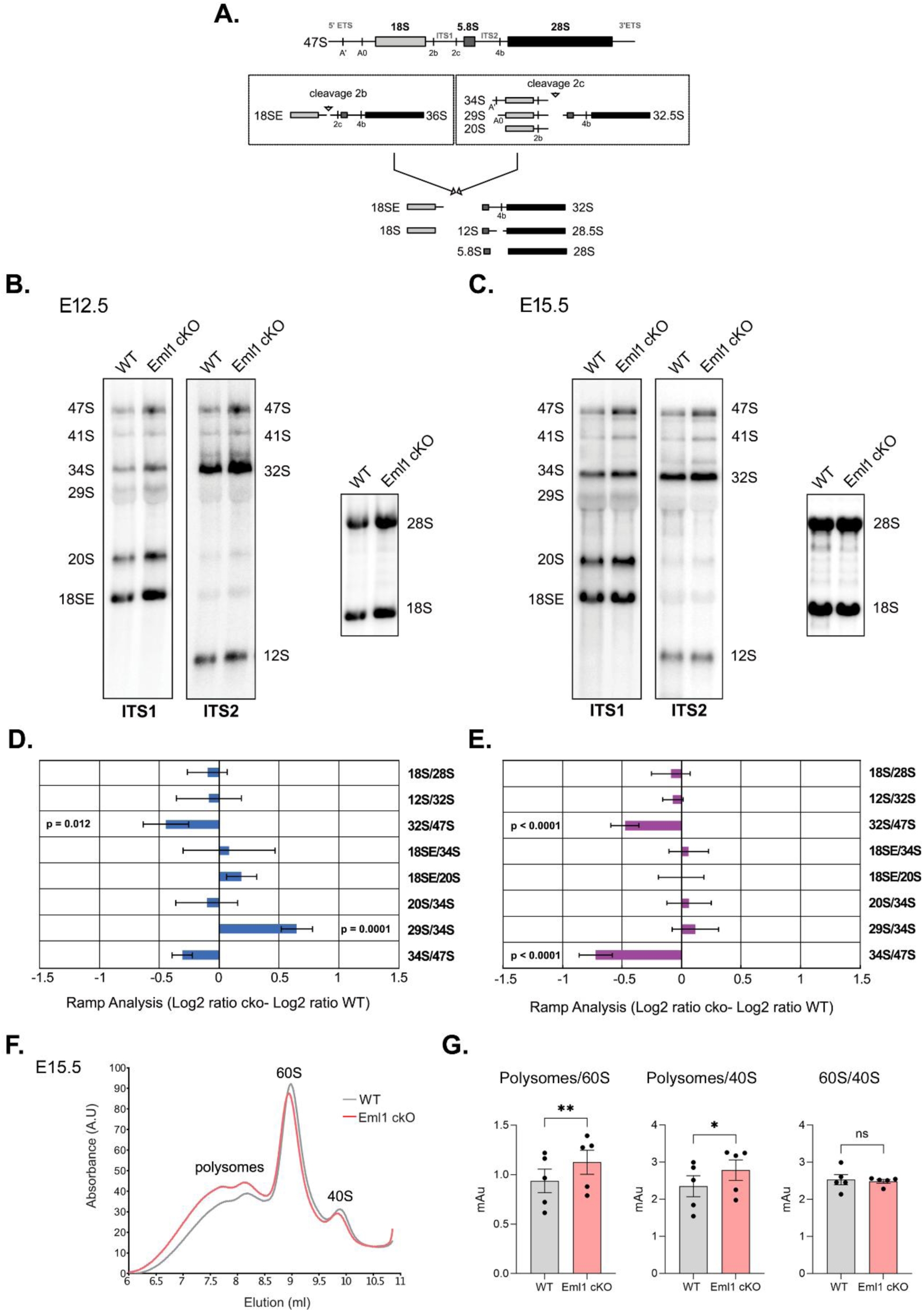
Increased ribosome biogenesis and polysomes in Eml1 cKO cortices. **A.** Pre-ribosomal RNA maturation process. **B, C.** Total RNA extracts prepared from WT and Eml1 cKO E12.5 (**B**) and E15.5 (**C**) cortices were analysed by northern blot. 3 µg of RNAs were separated on a 1.1% agarose gel and were probed with radiolabeled oligonucleotides complementary to ITS2, ITS1, or the 18S and 25S rRNAs (depicted in **A**). **D, E.** RAMP analysis of (**D**) E12.5 and (**E**) E15.5: the signals corresponding to the different rRNA precursors were quantified and log2 values of product to precursor ratios were expressed relative to control cells. N = 3 experiments. **F.** Polysome profile obtained with Ribo Mega-SEC performed on E15.5 cortices. The represented curve is a mean of the curves obtained from N = 5 experiments. **G.** Quantification of the ratio between polysomes and 60S, polysomes and 40S and 60S and 40S, showing an increase in polysomes in Eml1 cKO cortices at E15.5. Values are represented as mean ± SEM. N = 5 experiments from 5 different litters, 3 embryos were pooled per genotype. Paired t-test, p-value ** ≤0.01, * ≤0.05.

Ribo Mega-SEC polysome profiles can be analysed to obtain information concerning the ratio between isolated inactive ribosomal subunits and translationally active polysomes present in the samples. Comparing WT and Eml1 cKO profiles, we measured the area under the peaks and calculated the ratios for polysomes/60S, polysomes/40S and 60S/40S, at E12.5 (Supplementary figure 3B, C) and E15.5 (Figure 6F). While E12.5 analyses did not show differences in these ratios (Supplementary Figure 3C), E15.5 Eml1 cKO cortices showed a statistically significant increase in polysomes in Eml1 cKO samples (Figure 6G).

Together, these results suggest that the maturation of pre-ribosomal rRNAs is perturbed at both ages, suggesting especially at E15.5, mild ribosome synthesis increase. Ultimately, polysomes are increased in the Eml1 cKO at E15.5.

### Translating Ribosome Affinity Purification of WT and Eml1 cKO aRG suggests downregulation of translation in Eml1 cKO cells

We show that polysome distribution in endfeet is abnormal, and that ribosome biogenesis and polysomes are increased in the absence of Eml1. We hence decided to assess translation in Eml1 WT and cKO aRG.

Using first the Translating Ribosome Affinity Purification (TRAP) technique, translatomes of WT and Eml1 cKO aRG were assessed. TRAP is classically based on transgenic mouse lines expressing the chimeric ribosomal protein L10A tagged to GFP under the control of a cell-type specific promoter, allowing the purification of EGFP-tagged polysomes (Heiman et al. 2008, Doyle et al. 2008, Heiman et al. 2014). Variants of the technique that use IUE to target a specific cell type have also been described (Rannals et al. 2016, Huang et al. 2019). In the absence of a specific transgenic TRAP line for aRG, we decided to perform the latter technique to analyse these cells. Specifically targeting progenitors, E14.5 embryos were electroporated in utero with EGFP-Rpl10a. One day later, Rpl10a expression was detected in aRG cytoplasm and nucleoli, as expected for a ribosomal protein, as well as in apical and basal processes and endfeet (Figure 7A, Supplementary Figure 4A). Thus, after 24 h, E15.5 cortices were dissociated, and EGFP-labelled ribosomes were isolated by affinity purification. Actively translated mRNAs were harvested, and cDNA libraries were then sequenced (Figure 7B).

**Figure 7.**
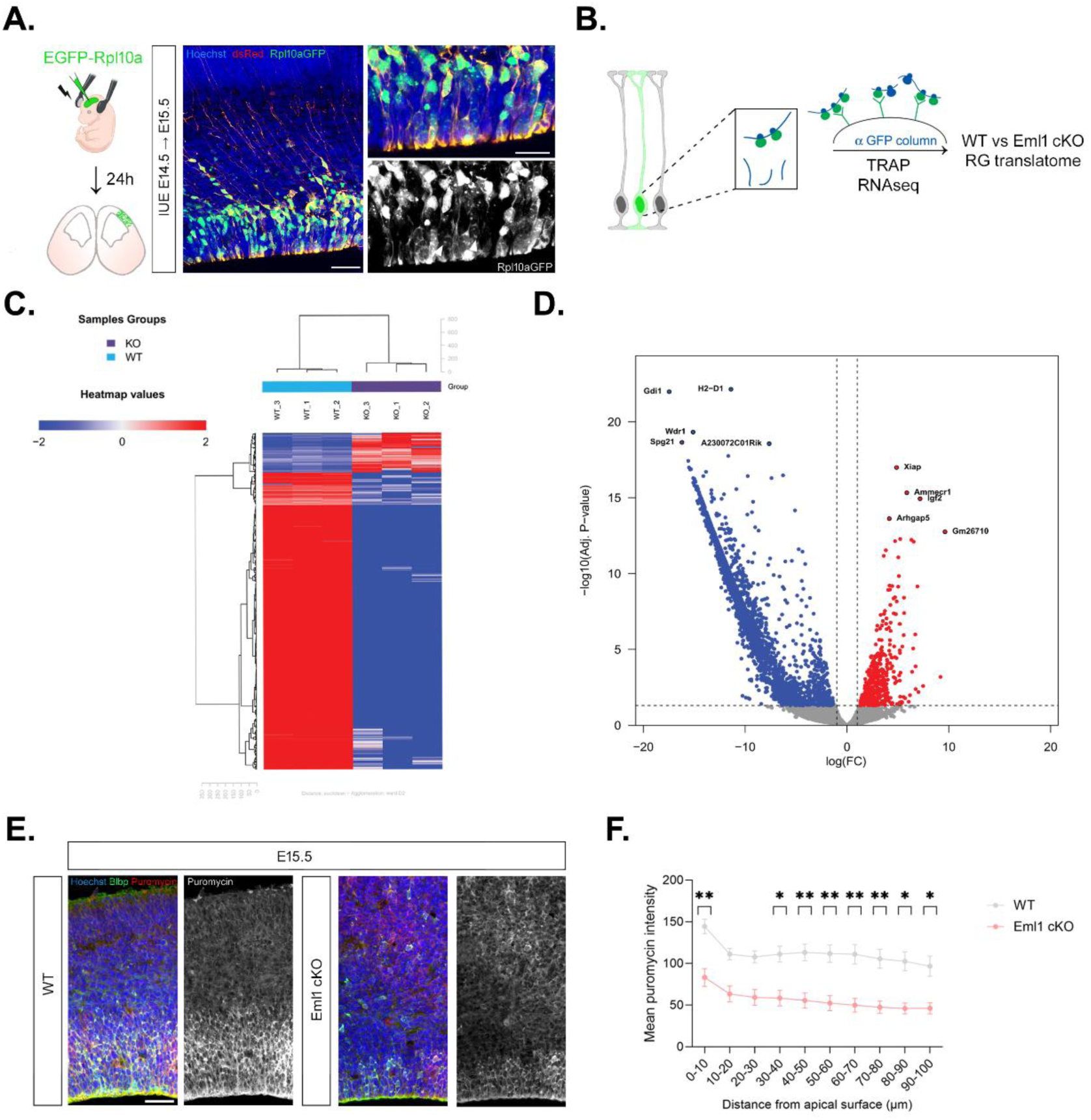
Translatome of WT and Eml1 cKO aRG. **A.** In utero electroporation of EGFP-L10a and dissection after 24 h allows the expression of the plasmid in aRG. Rpl10a is detected in the cytoplasm and nucleoli (arrows), as well as in apical endfeet and apical and basal process. Scale bar: 40 μm; magnifications: 20 μm. **B.** Workflow of TRAP experiments. GFP+ ribosomes were expressed in a subset of cells, allowing their isolation by affinity purification. Ribosome-bound mRNAs were then harvested and sequenced. **C, D.** Hierarchical clustering (**C**) and volcano plot (**D**) of differentially regulated genes in WT and Eml1 cKO aRG translatome. Fold change ≥ 2, adjusted p-value ≤ 0.05. N = 3 replicates per genotype from 3 different litters, each obtained by pooling at least 2 embryos. **E.** Puromycin staining in E15.5 cortices. **F.** Quantification of puromycin intensity from the apical surface. Values are represented as mean ± SEM. N = 8 (WT) and 6 (Eml1 cKO) embryos from 3 different litters. Mixed effect analysis, p-value ** ≤0.01, * ≤0.05. Scale bar, equivalent for WT and Eml1 cKO: 40 µm.

Bioinformatic analyses revealed a total of 4941 differentially translated genes between WT and cKO Eml1 cells, of which 4172 were downregulated (84.4%) and 769 were upregulated (15.6%) in Eml1 cKO (Figure 7C, D) (Table 3). A number of the genes identified were confirmed to be expressed in aRG (Telley et al. 2019, humous.org). Amongst dysregulated genes, we found classical aRG genes (e.g. *Pax6*, *Fabp7*, *Hes1*) as well as genes involved in progenitor proliferation (e.g. *H3f3b*, *Ybx21, Rbbp4*) (Xia and Jiao 2017, Dhanya et al. 2024, Zhang et al. 2025) and cell adhesion (e.g. *Cdc42*, *Arpc2, Actr3*) (Yokota et al. 2010, Wang et al. 2016).

**Table 3.**
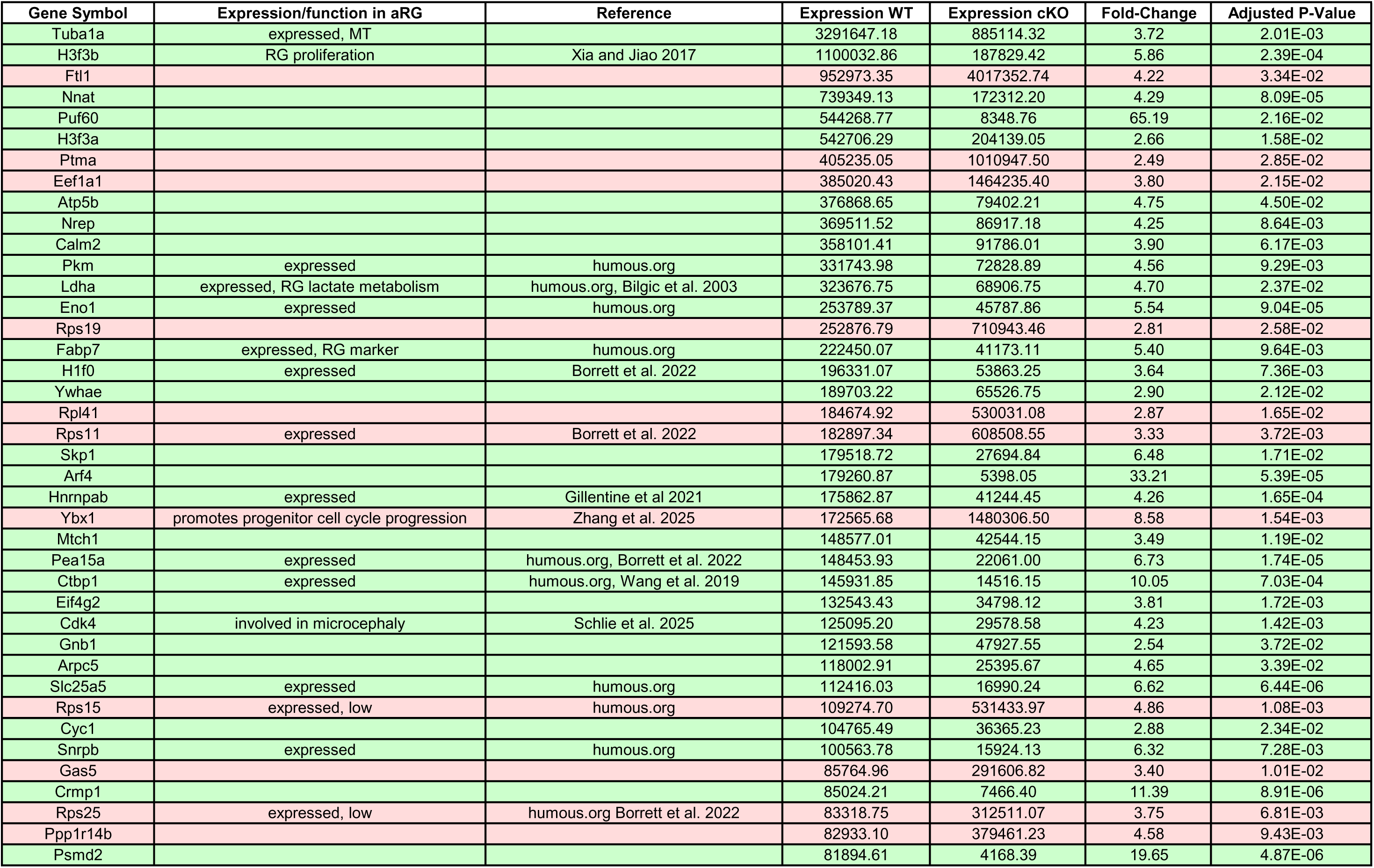
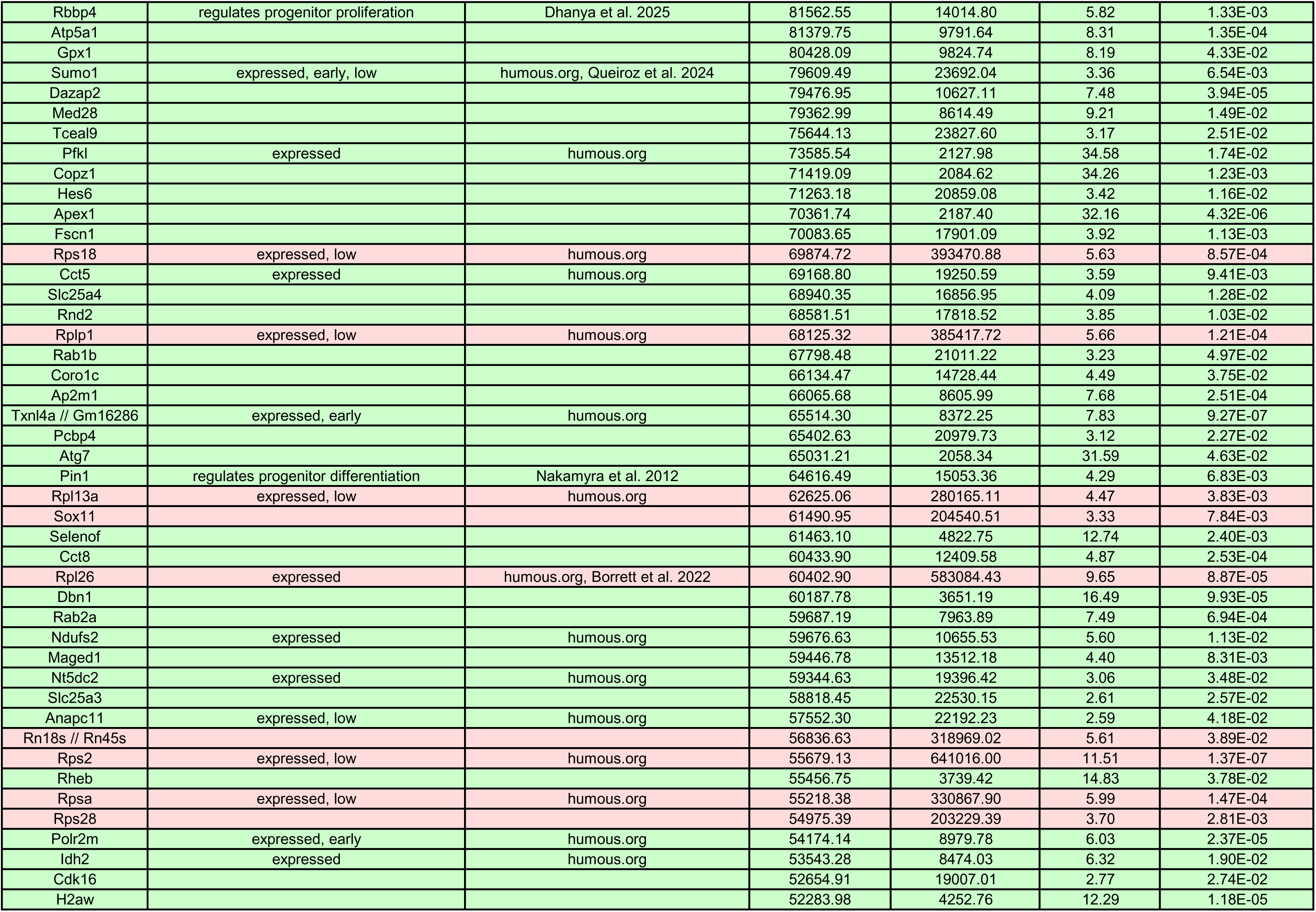
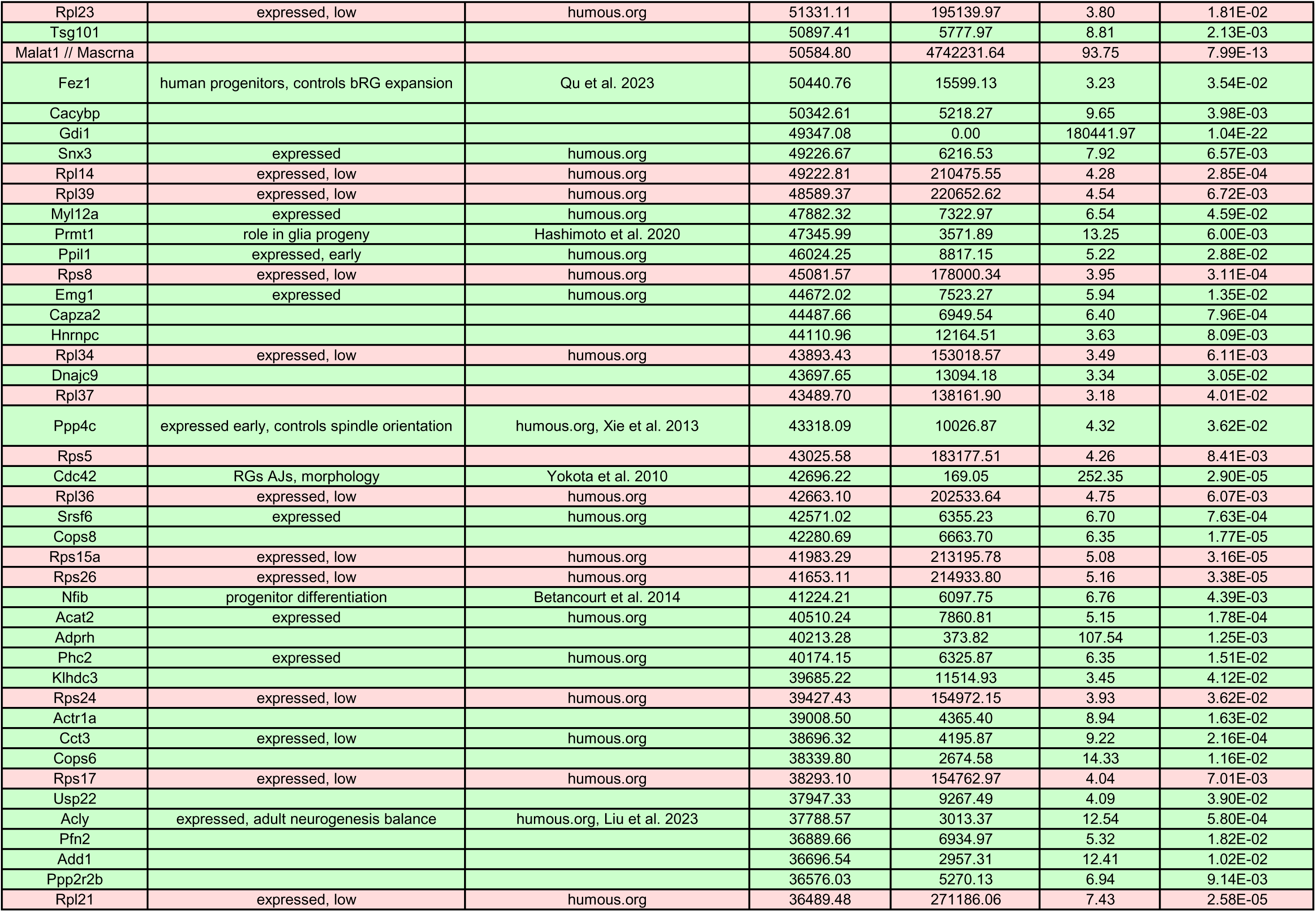
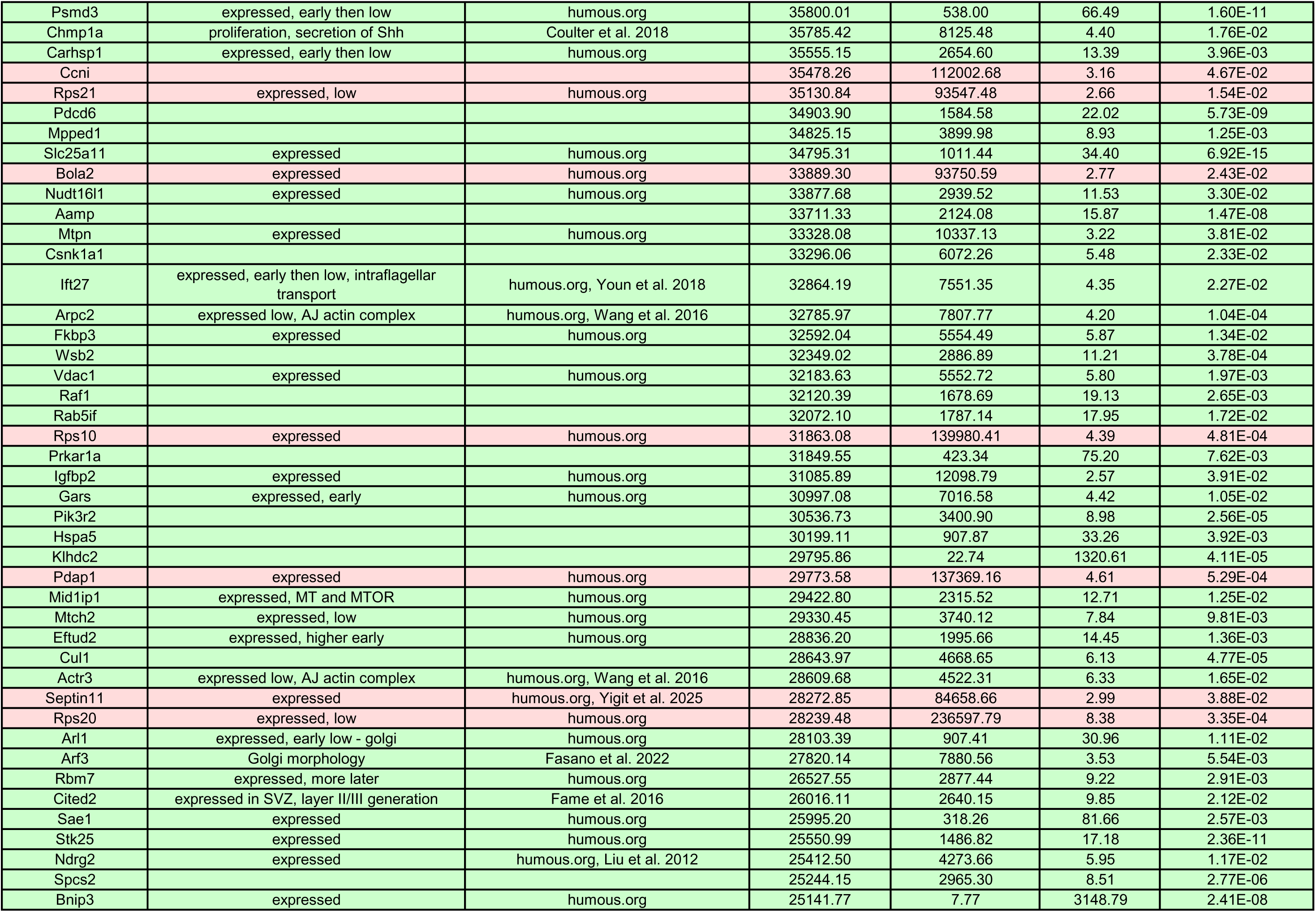
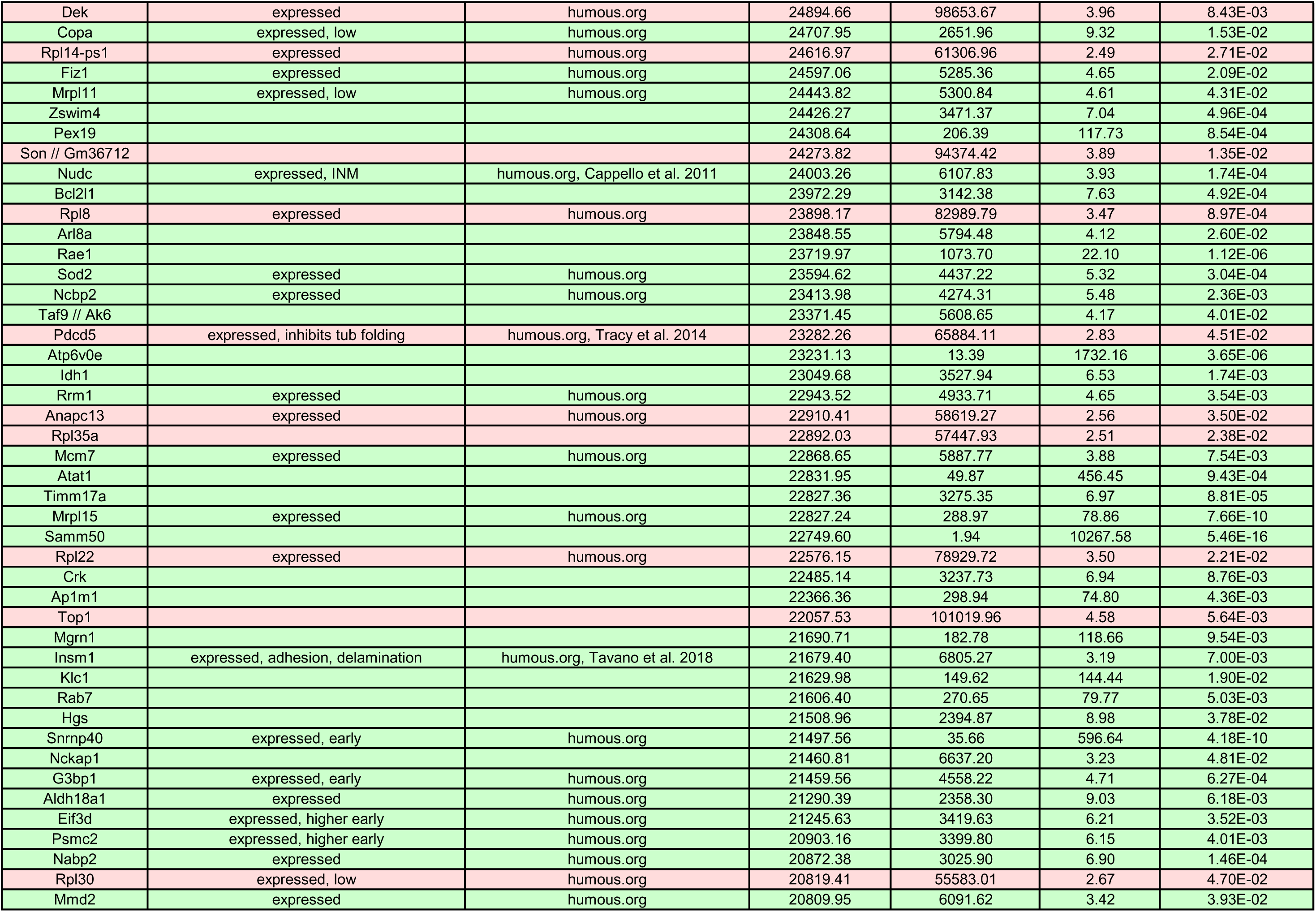
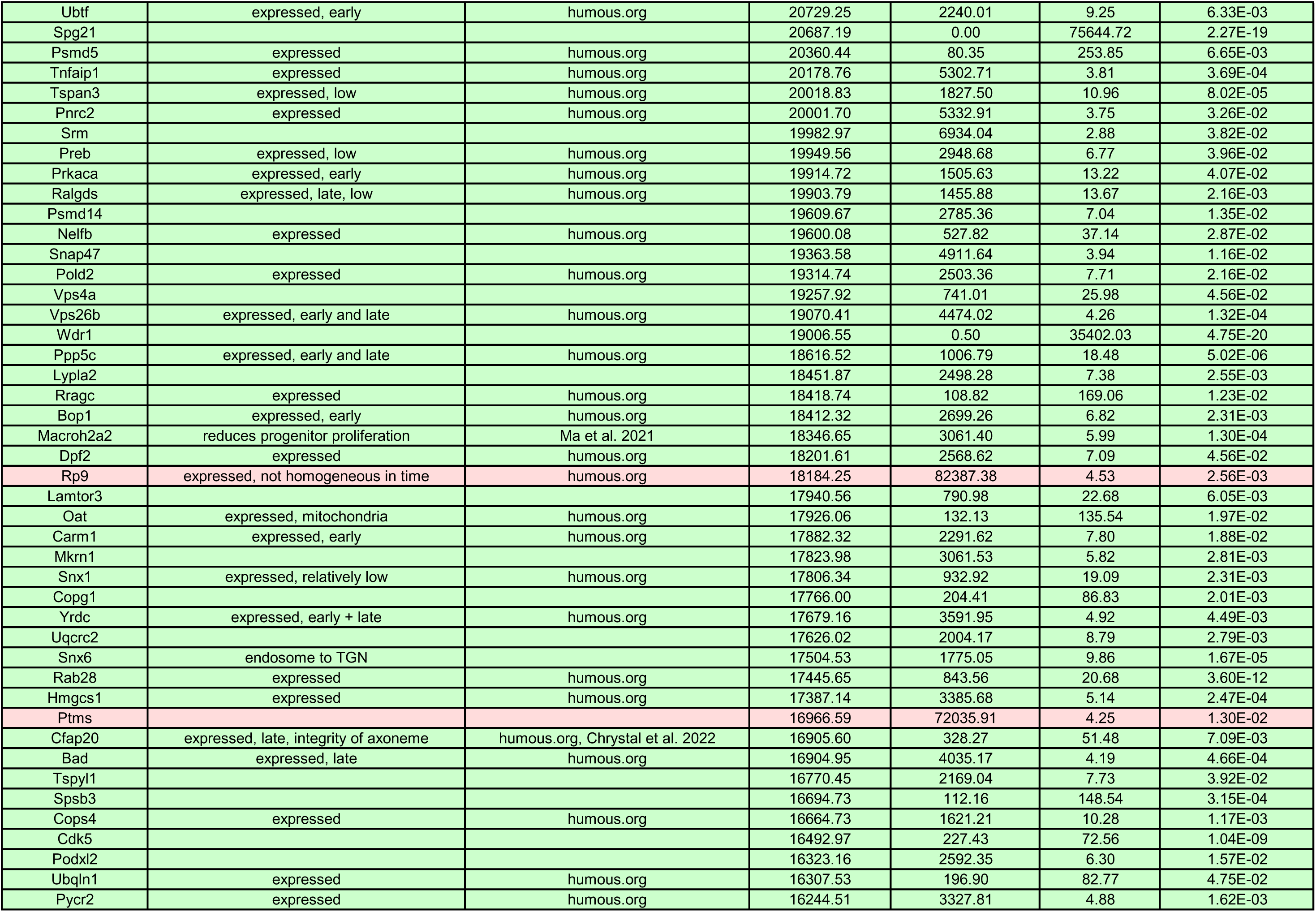
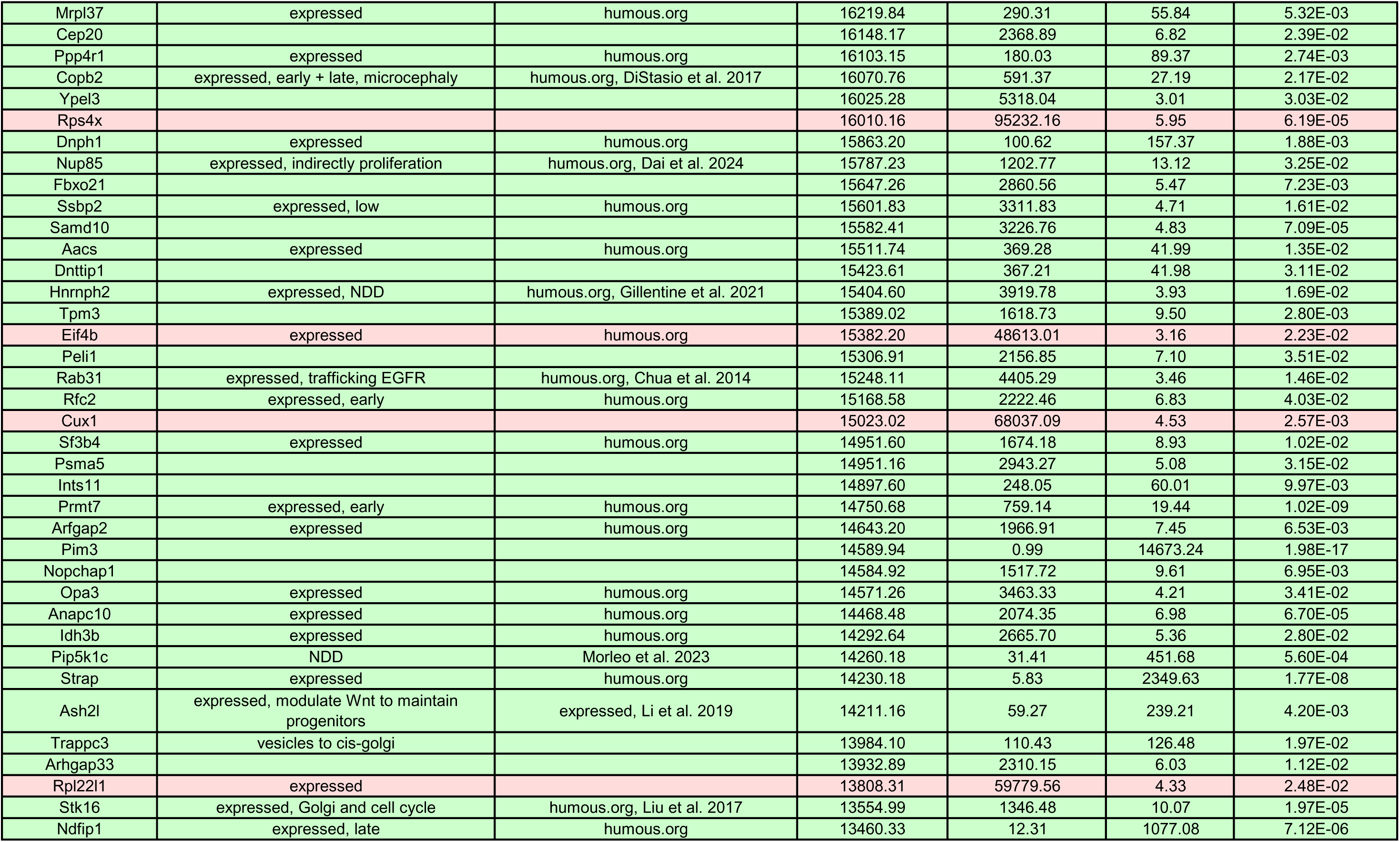
Expression and function in aRG of the 300 most expressed mRNAs in WT cells from TRAP experiments. Green: down in Eml1 cKO, red: up in Eml1 cKO

The TRAP results suggest an important downregulation of translated mRNAs in Eml1 cKO aRG. This was further validated by assessing levels of translation with puromycin injection in pregnant dams. Puromycin is an analogue of tRNA, it is therefore incorporated in nascent polypeptides and can be visualised by immunofluorescence (Figure 7E) (Chau et al. 2018, Enam et al. 2020). Puromycin incorporation indeed appeared lower in Eml1 cKO cortices at E15.5 compared to WT (Figure 7E, F), confirming the TRAP results. GO and Reactome analyses were performed to highlight the top significant terms and pathways amongst downregulated genes identified in the translatome data. GO terms related to the cytoskeleton and transport, as well as glucose metabolism, linked to mitochondria, were identified, supporting the phenotypes previously described (Supplementary Figure 5B, C). In the Reactome pathway analysis, mTORC1-mediated signalling appears as a significant pathway (45^th^ on 166 pathways identified). The mTOR pathway is a key regulator of protein synthesis, both in a general way and by controlling translation of specific mRNAs (Yang et al. 2022). Several genes of mTORC1-mediated signalling are downregulated in Eml1 cKO aRG, such as *mtor*, *lamtor3* and *rptor* (Supplementary Figure 5D), possibly at the origin of overall translation downregulation.

Amongst specific genes translated in WT aRG, and whose translation is downregulated in Eml1 cKO aRG, *Tuba1a* was identified (3.72-fold change). As previously mentioned, ECL sandwich immunoassays revealed a tendency for decreased tubulin in Eml1 cKO cell pellets and E12.5 cortices (Supplementary Figure 1C, D). α-tubulin immunostaining at E13.5 similarly revealed a tendency for reduced protein expression in the first 100 μm from the ventricular surface (Supplementary Figure 5E-G). Therefore, the translatome results validate previous findings (Zaidi et al. 2024, Yigit et al. 2025) and support the idea that Eml1 mutation impairs the MT cytoskeleton, including through translational control.

### Upregulated genes in Eml1 cKO aRG translatome are related to ribosomes and translational machinery

The translatome results are in contrast with the increase in ribosome biogenesis and polysomes previously described. As for the downregulated genes, GO and Reactome analyses were performed to highlight the top significant terms and pathways amongst upregulated genes (Figure 8). Amongst GO biological process (BP), cellular component (CC), molecular function (MF) and Reactome, the most enriched terms or pathways were found to be associated with ribosomes or translation, such as the terms “cytoplasmic translation” (BP) (Figure 8A), “polysome” and “ribosome” (CC) (Figure 8B), “structural constituent of the ribosome” (MF) (Figure 8C) and pathways related to 40S subunits and translation initiation complexes (Figure 8D). Both Rps and Rpl mRNAs are identified, as well as translation initiation factors. Interestingly, the genes of these categories were mostly upregulated (Figure 8E, F). The mild increase in ribosome biogenesis and polysomes detected by RiboMegaSec could be reflected by these upregulated genes.

**Figure 8.**
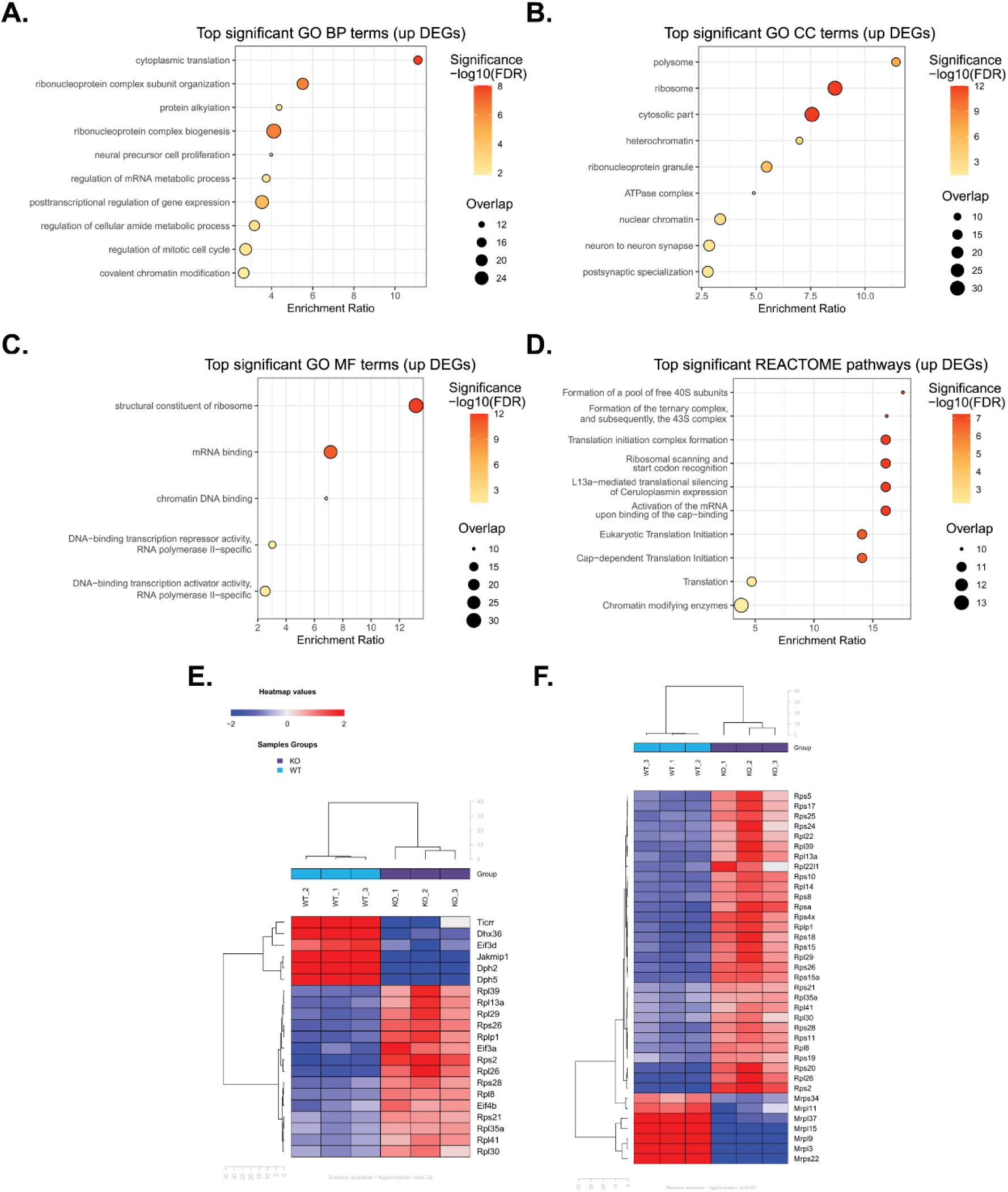
Top significant gene ontology terms and pathways indicate upregulation of translation-related genes. A-C. Top significant GO terms for (**A**) BP, (**B**) CC and (**C**) MF of upregulated genes. **D.** Top 10 significant Reactome pathways of upregulated genes. **E, F.** Hierarchical clustering of deregulated genes from the most enriched GO terms of (**E**) BP “cytoplasmic translation” and (**F**) MF “structural constituent of ribosome”. BP, biological process; CC, cellular component; MF, molecular function.

The most upregulated mRNA in Eml1 cKO aRG is *Rps2*. By immunohistochemistry we observe a tendency for increased protein expression at the ventricular surface of E13.5 cortices (Supplementary Figure 5H, I). In addition, ribosomal proteins are upregulated in proteomic screens (Yigit et al. 2025).

These results, together with tubulin immunostaining, suggest that certain differences observed in the translated mRNAs are reflected also at the protein level.

Overall, our data suggest a downregulation of translation (translatome and puromycin) in Eml1 mutant cells, despite an upregulation of ribosome and translation machinery genes, possibly as a compensatory mechanism. Reduced overall translation, and changes at apical and basal endfeet, contribute to an aRG phenotype associated with SH.

## Discussion

In this study, we describe cellular and subcellular defects of Eml1 cKO aRG, highlighting a new role for Eml1 in relation with organelle trafficking and function. More specifically, we describe that 1) mitochondria and polysomes are abnormally distributed in Eml1 cKO aRG, potentially due to trafficking defects; 2) Eml1 interacts with ribosome and translational machinery components; 3) overall translation levels are downregulated in Eml1 cKO aRG and 4) amongst upregulated translated mRNAs, many are related to translational machinery. These novel results raise several questions.

### Mitochondria trafficking and distribution

Eml1 was shown to alter general MT-based trafficking in Eml1 cKO Pax6+ progenitors in culture (Uzquiano et al. 2019, Zaidi et al. 2024). Also, recent proteomics data highlighted downregulation of proteins involved in organelle transport (Yigit et al. 2025). We demonstrate that mitochondria in Eml1 cKO progenitors have increased speed and accordingly decreased pausing time, particularly in the somal direction in cell processes in dissociated cells in culture. While outward trafficking depends on kinesin motor proteins, inward trafficking to the minus end of MTs is mostly driven by dynein, in association with several known adaptors such as TRAK for mitochondria (Sheng 2014, Reck-Peterson et al. 2018). Whether the MitoTracker results are related to a consequence of increased dynein/decreased kinesin action remains to be seen. Nevertheless, dysregulation of dynein subunits and kinesins was observed in omics data, as highlighted in Table 2.

Mitochondria dynamics and metabolism are already known to be important for neurogenesis (Iwata 2020, Iwata and Vanderhaeghen 2021). In aRG, they are distributed throughout the cell, in the soma, processes and endfeet (Rash et al. 2018). By EM, we show that in WT they are particularly enriched in basal endfeet at E12.5, but less so at E15.5. In Eml1 cKO brains, we detected an increased density in the apical endfeet and decreased density in the basal endfeet at both stages. The presence of mitochondria in distal apical and basal endfeet is likely to be linked to local functions, such as energy production and regulation of signalling (Rash et al. 2018, Cioni et al. 2019).

MitoTracker experiments were performed in vitro, where the centrosomes and therefore the minus ends of MTs are perinuclear. In vivo, they localize in the aRG apical endfeet (Wimmer and Baffet 2023). Thus, if increased ‘inward’ movement is also observed in vivo, this would reflect increased apical-directed transport of mitochondria, potentially leading to their accumulation, as observed in apical endfeet by EM. It is hence important to confirm trafficking alterations in vivo, where the apico-basal polarity is preserved as well as the local environments.

### Polysome distribution, interaction with Eml1 and trafficking

We also looked at polysome distribution and density across corticogenesis in Eml1 cKO aRG. Equal densities at E12.5, and relatively fewer in basal endfeet at E15.5 were observed in WT aRG. Similarly to mitochondria, Eml1 cKO aRG showed, at both stages, an increased density in the apical endfeet and decreased density in the basal endfeet. This is intriguing and likely to impact function, as local translation is known to take place in aRG basal endfeet (Pilaz et al. 2016).

We highlight the interaction of Eml1 with ribosomal proteins, as well as ribosome biogenesis and translation factors, across multiple interactome analyses (Bizzotto et al. 2017, Uzquiano et al. 2019, Zaidi et al. 2024). Eml1 also associated with 40S ribosomal subunit fractions in E15.5 cortices. Rpl proteins identified in interactome experiments could interact indirectly, or have transient interactions, with Eml1, as no obvious direct association with the 60S subunit was observed. The Eml1 ortholog EMAP was suggested to link ribosomes to MTs in Sea urchin egg MT preparations (Suprenant et al. 1993, Hamill et al. 1994). Due to the high conservation of the proteins of this family (Suprenant et al. 2000, Richards et al. 2014), it is possible that Eml1 could share this function of binding to ribosomes, maybe particularly with the 40S subunit.

Little is known about the specific interaction between ribosomes/translational machinery and MTs. Several studies have nevertheless shown polysomes in association with or in proximity of MTs (Suprenant 1993, Han et al. 2006, Mahamid et al. 2016). In addition, proteins that co-precipitate with MTs after MT pelleting always include ribosomal proteins, translation initiation and elongation factors, (Sakamoto et al. 2008, reviewed in Chudinova et al. 2018). The translation initiation factor eIF3a also associates with MTs (Shanina et al. 2001, Sakamoto et al. 2008). In addition, the neuronal MAP Tau binds to Rps6 (Meier et al. 2016). These results suggest that MAPs, and more widely the MT cytoskeleton, bind ribosomal proteins and translational machinery components, and this may be an overlooked area of study.

Given our combined results, we propose that Eml1 could be involved in ribosome and/or polysome transport or attachment to MTs. Eml1 is not an MT motor itself, thus it may act as an adaptor protein between kinesin/dynein motors and the ribosome/polysome cargo (Bartoli et al. 2011, Chudinova et al. 2019). Several kinesins were identified as potential interactors of Eml1 in pulldown experiments, such as Kif22 (Bizzotto et al. 2017), and other Eml1-Kif interactions are likely. However, we also can point out that as Eml1 can itself bind to and influence MTs, it may also be involved in ribosome/polysome attachment.

Not much is known about ribosome and/or polysome transport but it is reasonable to assume that ribosomes are transported to the different cell compartments where they exert a local function (Pilaz and Silver 2017, Mazaré et al. 2021, Broix et al. 2021). Scattered evidence across cell types, including neurons, support the role of kinesin motors in the transport of the translational machinery (Sotelo-Silveira et al. 2004, Bisbal et al. 2009, Graber et al. 2013, Song et al. 2015, Scarborough et al. 2021). Defects in trafficking in Eml1 cKO progenitors could explain the accumulation of polysomes in apical regions, close to the site of ribosome biogenesis (the nucleus and nucleolus), since long-distance trafficking is required to reach the more distant basal endfoot.

### Reduced MT stability in Eml1 cKO

It is possible that the defects in organelle distribution are linked to overall MT instability, rather than trafficking defects. Several studies using different models have highlighted such mechanisms, showing that MT disruption caused by mutations or drug treatment leads to perturbed ribosome as well as mitochondria distribution (Noma et al. 2017, Heo et al. 2018, Denes et al. 2021, Scarborough et al. 2021). This suggests a conserved role of MT stability in organelle distribution in the cells.

*Eml1* mutation impacts MT nucleation and growth (Bizzotto et al. 2017, Zaidi et al. 2024). Here, we also show that *Tuba1a* translation is downregulated in Eml1 aRG and ECL-based immunoassays also revealed a tendency for decreased tubulin in Eml1 cKO cell pellets and E12.5 cortices. Tubulin was also found to be decreased in Eml1 cKO pellets (Yigit et al. 2025) and its fluorescence intensity is decreased at E12.5 and E13.5, particularly at the ventricular surface of Eml1 cKO cortices (Zaidi et al. 2024), as well as in Eml1 knockdown HeLa cells (Yigit et al. 2025). Thus, there are multiple results suggesting subtle MT alterations. It is still unclear however, how these might impact trafficking and alter distribution.

We explored the status of tubulin PTMs, known to regulate MT functions (Janke and Magiera 2020). ECL-based immunoassays of key tubulin PTMs did not show any major differences, even though some tendencies were observed. Eml1 may still play a role, based on studies in neurons where Eml1 absence impaired axonal growth and led to reduced acetylated tubulin, rescued by treatment with deacetylase inhibitors (Zhang et al. 2024). The authors also show interaction between Eml1 and αTubulin acetyltransferase 1 (αTAT1) (Zhang et al. 2024). EML2, a member of the EML family, preferentially binds tyrosinated MTs (Hotta et al. 2022). Also, in aRG TRAP results we observe downregulation of glutamylases and acetylases (e.g. *Ttll5*, *Atat1*) as well as their counterparts deglutamylases and deacetylases (e.g. *Atgpbp1*, *Hdac6*). Thus we cannot rule subtly altered tubulin PTMs in progenitors.

### Alterations of endfeet structures in Eml1 cKO aRG

The apical and basal endfeet of aRG are highly specialized compartments with different structures and organelle compositions (Viola et al. 2024). In addition, they are exposed to diverse cues, as the apical side faces the ventricle and the basal is in contact with the meninges (Ferent et al. 2020). Their integrity is crucial for cell polarity and function. We speculate that the abnormal densities of polysome and mitochondria in apical and basal endfeet of Eml1 cKO aRG could alter their morphologies and functions. We focused on adhesive contacts, endfoot size and overall aspect.

Loosening or disruption of cell AJ have been previously associated with both physiological and pathological progenitor cell delamination (Cappello et al. 2006, Jossin et al. 2017, Tavano et al. 2018, Camargo-Ortega et al. 2019). AJ length is reduced in Eml1 cKO compared to WT at E12.5 but not E15.5 and this is coherent with the proposed time-frame of delamination (Zaidi et al, 2024).

Apical extremity size is also increased in Eml1 cKO brains. It has been described that defects in centrosomes and PC lead to apical endfeet enlargement (Foerster et al. 2017, Shao et al. 2020). In *Eml1* mutant mouse models, both PC and centrosome defects are observed, and this phenotype is more severe at early stages (Uzquiano et al. 2017, Zaidi et al. 2024). It is currently unknown if increased size may be linked to an increased number of apical organelles and if this phenomenon is also observed at later stages e.g., at E15.5. In addition, restoring the MT cytoskeleton by the use of EpoD might reset the organelle distribution and consequently reduce apical endfeet size.

A reduced number of basal radial fibers was observed in Eml1 cKO cortices at E15.5, and the remaining basal endfeet also appear simpler. *Marcks* deletion in aRG leads to perturbed and reduced fibers spanning the cortex and simpler basal endfeet compared to WT, probably due to the role of Marcks in anchoring components of signalling complexes (Weimer et al. 2009). *Myh9* cKO brains also display reduced basal endfeet per cell, with fewer ramifications, probably linked to the cytoskeletal regulation role of Myh9 (D’Arcy et al. 2023). Therefore the basal phenotypes we observe might be linked to Eml1 MT function, ultimately controlling cell shape and organization, likely to be influenced by organelle distribution.

### Translation downregulation in Eml1 cKO aRG

TRAP experiments were performed by IUE of Rpl10a-GFP in aRG. GFP+ ribosomes were isolated from WT and Eml1 cKO aRG and their associated mRNAs sequenced. Interestingly, 84.4% of the differentially translated genes identified were translationally downregulated in Eml1 cKO, and puromycin staining of the embryonic cortices at E15.5 confirmed this result. It is interesting that MT cytoskeletal components amongst others appear impacted. As an example, *Tuba1a* mRNA appears downregulated and similar tendencies were observed at the protein level by immunostaining.

In the absence of Eml1, translation is hence greatly reduced. We described Eml1 association in 40S fractions. The 40S subunit is at the core of the translation initiation machinery (Brito Querido et al. 2024). It is hence possible that Eml1 plays a role in translation initiation and that in its absence, defects in the initiation steps would lead to overall translation downregulation. Furthermore, Ddx3x, whose mutations are known to underlie neurodevelopmental disorders, has been shown to play an important role in translational control at early stages of corticogenesis, as its absence reduced the translation efficiency of several transcripts (Hoye et al. 2022). Indeed, our BioID experiments suggest an interaction between Eml1 and Ddx3x. In addition to binding ribosomes as previously mentioned, Tau has also been shown to play roles in translation (Meier et al. 2016, Evans et al. 2019, Banerjee et al. 2020). This suggests that there might be conserved roles of some MT-binding proteins regulating translation and this understudied phenomenon may be used by the cell to regulate key processes.

### A potential compensation mechanism increasing the level of translation machinery

In the top significant GO terms and pathways of upregulated genes, we identify, paradoxically, ribosomes, polysomes and translation, and particularly translation initiation. Rps4x, Rps10 and Rps20 are also upregulated in proteomic data (Yigit et al. 2025). In Ribo-Mega-SEC experiments, we observed an increase in polysomes at E15.5 and Northern blot experiments using RNA extracted from the same samples suggest a mild increase in RNA polymerase I activity, strongest at E15.5. It is therefore possible that the cells attempt to restore translation levels by boosting initiation and by producing more ribosomes. Studies of translational control across corticogenesis have shown translation downregulation at E15.5 that targets the translation machinery itself, associated with an acute decrease in ribosome biogenesis (Harnett et al. 2022). Eml1 in WT conditions might play a role in this process, since we observe that its absence leads to an opposite phenotype.

These results are in contrast with the overall translation downregulation suggested by TRAP and puromycin experiments. Ribo-Mega-SEC experiments were however, performed on whole cortex lysates containing aRG, IP and neurons, we cannot therefore exclude that the increased polysomes could be due to other cell types than aRG. On the other hand, ribosome biogenesis factors were identified as potential Eml1 interactors (e.g. Fbl, Nat10, Ncl) and the function of Eml1 related to ribosome biogenesis remains worth investigating. Of interest, Tau localizes to fibrillar components of the nucleoli, where ribosome biogenesis takes place (Loomis et al. 1990).

Cell surveillance mechanisms detect defective mRNA translation, to restore ribosome homeostasis (Mills and Green 2017). In Epstein-Barr virus infections, inhibition of translation leads to an increased ribosome biogenesis and cell proliferation (reviewed in Vadivel Gnanasundram and Fahraeus 2018) and division into two daughter cells involves temporary inhibition of translation (Pyronnet and Sonenberg 2001, Sivan and Eroy-Stein 2008). During corticogenesis, aRG cell cycle length has been linked with differentiation (Calegari and Huttner 2003, Hardwick et al. 2015), and Eml1 cKO E15.5 cells have a reduced cell cycle length compared to WT and increased proportion of Pax6+ cells (Zaidi et al. 2024). Thus, in E15.5 Eml1 cKO cells, we observe increased proliferation compared to WT (Zaidi et al, 2024), reduced translation, and signs of increased ribosome biogenesis. It appears hence that there may be interdependent dysregulated translational control and cell proliferation in the Eml1 cKO. Thus, we pinpoint several understudied interdependent processes, involving MTs, translation and aRG function during neurodevelopment, to be further explored in the future, as they appear crucial for regulating these key progenitors during cortical development.

## Materials and methods

### Animals

Research was carried out conforming to national and international directives (directive CE 2010/63 / EU, French national APAFIS n° 23424; 46509) with protocols followed and approved by the local ethical committee (Charles Darwin, Paris, France). Mice were housed with a light/dark cycle of 12 h (lights on at 07:00). Males and females were used in all analyses. The Eml1 mutant line was generated and analyzed on the mouse genetic background C57BL/6N (B6N) (Zaidi et al, 2024). All mice were housed in the IFM Institute animal facility or at the CDTA, Orléans, France.

### Crosses and genotyping

Eml1 flox/flox animals were crossed with Eml1 flox/flox-Emx1Cre/+ animals. Females were placed in the male cage and the following morning the presence of a vaginal plug was observed and considered to be embryonic day 0.5 (E0.5). Embryonic brains were collected at the indicated times. Genotyping primers used to detect Cre were:

Cre 1: 5’-GAA CCT GAT GGA CAT GTT CAG G-3’.

Cre 2: 5’-AGT GCG TTC GAA CGC TAG AGC CTG T-3’ Cre3: 5’ -TTA CGT CCA TCG TGG ACA GC-3’

Cre4: 5’ -TGG GCT GGG TGT TAG CCT TA-3’

Primers used to detect the floxed Eml1 allele were:

Primer Lf: 5’-GAA AAC GTG CTT TGC TGT GTA CAT AGG-3’.

Primer Er2: 5’-CTT GTT AAA GCG TCT GCA GTC TGT CTG-3’.

### Antibodies and plasmids

Primary antibodies used were: rabbit anti-Blbp (Sigma PRS4259), rabbit anti-Eml1 C3 (GeneTex GTX100252, 1:300), mouse anti-Fibrillarin (Abcam AB4566, 1:500), rabbit anti-GFP (Invitrogen A6455; 1:1000), mouse anti-Puromycin (Millipore AF647, 1:2000), guinea pig anti-RFP (Synaptic systems 390004, 1:1000), mouse anti-Rpl5 (non-commercial), mouse anti-Rps15 (non-commercial), rabbit anti-Rps2 (GeneTex GTX131192, 1:200), mouse anti-α-tubulin (Sigma-Aldrich T9026, 1:500), mouse anti-γ-tubulin (Sigma-Aldrich T6557, 1:400). Secondary antibodies used were: anti-rabbit HRP (Promega W401B), goat anti-rabbit Alexa 488 (Thermo Fisher Scientific A-11008, 1:1000), goat anti-rabbit Alexa 633 (Thermo Fisher Scientific A-21070, 1:1000), goat anti-mouse Alexa 488 (Thermo Fisher Scientific A28175, 1:1000), goat anti-mouse Alexa 633 (Thermo Fisher Scientific A-21050, 1:1000), donkey anti-guinea pig (Jackson 706-166-148, 1:1000).

Detection antibodies for ECL-based immunoassay were rabbit anti-polyE (AdipoGen AG-25B-0030), rabbit anti-detyrosinated α-tubulin (RevMab Biosciences 31-1335-00), mouse anti-tyrosinated α-tubulin (Sigma-Aldrich T9028), mouse anti-acetylated α-tubulin (Sigma Aldrich T6793). SulfoTAG labelled secondary antibodies were anti-mouse (MSD, R32AC-5) and anti-rabbit (MSD, R32AB-1).

Plasmids used were Blbp-GFP (Kielar et al. 2014), pCAG-EGFP-Rpl10a (Rannals et al. 2014) and pCAG-dsRed (Addgene #11151) for IUE.

### Mouse neuronal progenitor primary cell culture

The neuronal progenitor cell cultures were adapted from Sun et al. (2011) giving a highly enriched population of Pax6(+) cells. For this, 6-well cell culture plates were coated with poly-D lysine (PDL, P6407; Sigma-Aldrich) 2 µg/cm2, 1 h at room temperature. PDL was then removed, the plates were rinsed once with sterile water and coated with 1 µg/cm2 fibronectin (F1141; Sigma-Aldrich) in sterile 1X PBS. E14.5 timed-pregnant mice were sacrificed by cervical dislocation and the uterus was placed in ice-cold basal medium (DMEM/F12 Hams, 21041; Thermo Fisher Scientific, 1% Pen-Strep [Gibco], 2.9 mg/ml glucose and 1.2 mg/ml sodium bicarbonate). The embryos were collected and the cortex from both hemispheres was dissected and kept at 4°C in basal medium. The medium was removed and substituted by prewarmed sterile complete medium (basal medium complemented with 1X B27 without vitamin A (12589-010; Gibco), 20 ng/ml of EGF (E9644; Sigma-Aldrich), and 20 ng/ml of FGF (F0291; Sigma-Aldrich). The tissue was dissociated, and each sample was centrifuged (5 min, 1,200 rcf). The medium was removed and substituted by fresh prewarmed complete medium followed by resuspension of the cells. 1 × 10^5^ cells were plated in coated six-well culture plates. The cells were split once at 7 days in vitro (DIV) before performing experiments. Half of the culture medium was changed by a fresh complete medium every 2 days for 1 wk. For splitting, cells were washed with prewarmed Versene (Gibco), followed by a 3 min incubation with prewarmed StemPro Accutase (Gibco) at 37°C. Cells were plated (6–8 × 10^5^) on coated 14-mm glass coverslips and cultured for 2 DIV for immunocytochemistry experiments.

### Live imaging of mitochondria in Pax6+ progenitor cells in culture

Pax6+ progenitor cells were cultured at a confluence of ∼70% with DMEM F12 medium. Cells were then exposed to MitoTracker 0.2 µM Deep Red FM (Thermo Fisher, catalog no. M22426) for 30 min at 37 °C before recording. The microscope chamber was heated to 37°C for live recording. Live imaging was performed using an inverted Spinning Disk Leica microscope at 100-ms frame intervals. Kymographs were generated for single-blind analysis using ImageJ plugin-KymoToolBox. Mitochondria were considered stationary at <0.1 μm/s.

### Immunocytochemistry

Cells were washed in 1X PBS prior to fixation with 4% wt/vol PFA in 0.1 M phosphate buffer, pH 7.4, for 15 min at RT or fixed with methanol at −20°C. The cells were extensively washed for 15 min in PBST (Triton X-100 0.1% in 1X PBS). Incubation with blocking solution (10% NGS; Thermo Fisher Scientific, 0.1% Triton X-100 in 1X PBS) was performed for 1 h at RT and primary antibodies were applied overnight (O/N) at 4°C (see above for antibodies). The cells were extensively washed with a blocking solution and secondary antibodies were incubated for 2 h at RT in the dark. After washes, Hoechst (1:10,000; Thermo Fisher Scientific) was applied for 15 min at RT in the dark. The cells were extensively washed in PBS and the coverslips were mounted with Fluoromount G (Southern Biotechnology). Images were acquired at room temperature with a TCS Leica SP5-II confocal microscope.

### Electron microscopy

Mouse embryos were transcardially perfused with a solution containing 4% PFA (Electron Microscopy Science), 2.5% glutaraldehyde in sodium phosphate buffer (PB) 0.1 M, pH 7.4. 3 h after perfusion, brains were dissected and post-fixed in the same solution O/N at 4°C. The samples were washed extensively with PB 0.1M, and afterward 3x 10 min washes with Palade buffer (Palade, 1952) were performed. The samples were incubated in 2% osmium tetroxide in Palade buffer for 40 min at RT, and then rinsed in Palade buffer for 3 min, followed by another 3 min wash in distilled water. They were dehydrated in a series of ethanol baths and flat-embedded in epoxy resin (EPON 812, Polysciences). After polymerization, blocks containing the dorsal telencephalon were cut at 70 nm thickness using an ultramicrotome (Ultracut E Leica). Sections were cut with a diamond knife and picked up on formvar-coated 200 mesh nickel one slot grids. Sections were examined in a Philips CM100 electron microscope. Digital images were obtained with a CCD camera (Gatan Orius).

#### Electron tomography, segmentation and ribosome/polysome quantification within electron tomograms

Sections of resin embedded WT or HeCo mouse cortices were cut on a UCT ultramicrotome (Leica) with a nominal thickness of 80 nm, and picked up on 150-mesh formvar-carbon-coated copper grids (Electron Microscopy Science). Ten nanometer protein A-gold conjugates (Aurion) were added to the surface of each section to serve as fiducial markers for subsequent image alignment in IMOD<u>^61^</u>. Sections were stained with 5% uranyl acetate in 70% methanol and lead citrate. Tomographic data were acquired from the 80-nm resin sections using a JEOL JEM-2100 transmission electron microscope operating at 200 kV. Images were recorded at a nominal magnification of 10,000×, corresponding to a calibrated pixel size of 9.74 Å at the specimen level. Tilt series were collected over a range of +60° to – 60° using 2° increments, with the tilt axis set to 0°. Tilt series alignment and 3D reconstruction were performed using the IMOD software package. Prior to reconstruction, the tilt images were binned by a factor of 2, producing a final pixel size of 19.48 Å in the reconstructed tomograms. Weighted back-projection was used for tomogram generation, and all tomograms were manually inspected for alignment quality, section integrity, and contrast uniformity. To improve the visibility of structural features and enhance the segmentation accuracy, NAD (Nonlinear Anisotropic Diffusion) filtering was applied to the tomograms prior to and during the segmentation process. Segmentation of ribosomes and polysomes was carried out using the TomoSeg module in EMAN2. Automated segmentations were subsequently reviewed and refined manually when necessary. Quantitative volumetric measurements of polysomes were performed in UCSF ChimeraX. Segmented polysomes were imported as volumetric datasets, and their volumes were computed using the “Measure Volume and Area” tool available in the Volume Data panel. All volumetric measurements were reported in ångströms (Å^2^), based on the calibrated voxel size of 19.48 Å corresponding to the reconstructed tomograms.

#### Tissue lysates

Embryonic cortices were collected, and lysis of each embryonic cortex was performed individually by resuspending the tissue continuously with lysis buffer for a period of 1 h at 4°C. The lysate was then centrifuged (30 min, 15,000 rcf, 4°C), the supernatant was collected, and the protein concentration was measured using the BCA protein assay kit (Thermo Fisher Scientific) and the BertholdTech Mithras ELISA microplate reader.

#### ECL-based immunoassays

Lysates of E12.5 cortices and Pax6+ cells from WT and Eml1 cKO mice were prepared at 0.5 mg/ml, and were further diluted to 0.03mg/ml. The 96-well plates (MSD, ref: L15XA) were coated with 25µl of sample (0.75µg total protein) overnight without shaking at 4°C. The next day, the wells were washed 3 times with 0.05% Tween-20 (Sigma) in 1X PBS (Gibco). Blocking was done with 5% Blocker A-1X phosphate buffer (150 µl/well) (MSD, R93AA) shaking at 700 rpm for 1 hour at room temperature. The wells were washed 3 times with 0.05% Tween-20 in 1X PBS to reduce non-specific signal. The samples were then incubated with the detection antibodies (25 µl/well) diluted in 1% Blocker A-1X phosphate buffer, shaking at 700 rpm for 2 hours at room temperature. The wells were washed 3 times with 0.05% Tween-20 in 1X PBS. The samples were then incubated with the sulfo-TAG-labeled antibody (25µl/well) diluted 1:2000 in 1% Blocker A-1X phosphate buffer, shaking at 700 rpm for 1 hour at room temperature. The wells were washed 3 times with 0.05% Tween-20 in 1X PBS, and the Gold Read Buffer A (R92TG-2), containing the ECL coreactant, was added to the wells (150 µl). The plate was read using the MSD QuickPlexSQ 120 MM instrument.

### Ribo Mega-SEC

#### Protein lysates and dosage

Five cortices for each genetic background were resuspended in 700 µL of extraction buffer (20 mM Hepes-NaOH (pH 7.4), 130 mM NaCl, 10 mM MgCl₂, 1% Triton X-100, 0.2 mg/mL heparin, 2.5 mM DTT, complete EDTA-free protease inhibitor, and 0.1 U/µL Rnasin). The cortices were lysed by sonication for 1 minute using 5-second on/off cycles at 20% amplitude on a Branson 250 sonicator. The resulting cell extracts were clarified by centrifugation at 16,000 × g for 10 minutes at 4°C and then filtered through a 0.45 µm filter (Millipore). Total RNA was quantified using a Nanodrop spectrophotometer at 260 nm. A 500 µL sample of RNA at 500 ng/µL was injected onto Ribomegasec columns, mounted in series on an ÄKTApure system. The column setup included a first Agilent Bio Sec-5 guard column (7.8 × 50 mm, 2000 Å), followed by a second Agilent Bio Sec-5 column (7.8 × 150 mm, 2000 Å), and terminated with a third Bio Sec-5 column (7.8 × 150 mm, 1000 Å). The columns were pre-equilibrated with equilibration buffer (20 mM Hepes-NaOH, pH 7.4; 60 mM NaCl; 10 mM MgCl₂; 0.3% Triton X-100; 0.2 mg/mL heparin; 2.5 mM DTT) before the RNA extracts were injected. The samples were separated at a flow rate of 0.3 mL/min, and elution profiles were recorded after RNA elution using a UV detector at 260 nm on the AKTApure system with Unicorn 6 software. The various peaks in the elution profiles were then analyzed and quantified using Fityk software.

#### Western blot

Fractions from ribomegasec were precipitated using the TCA method and resuspend in Invitrogen 2X sample buffer (NuPAGE™ LDS Sample Buffer (4X) (NP0007) and NuPAGE™ Sample Reducing Agent (10X) (NP0009) and loaded on NuPAGE™ 4 to 12%, Bis-Tris protein gels (Invitrogen, NP0321BOX) and transferred to nitrocellulose membrane using the Trans-Blot Turbo Transfer System from Biorad. Membranes were immunoblotted with a primary antibody, followed by incubation with a secondary antibody coupled with horse radish peroxidase (HRP, Promega anti-rabbit (W401B)). The blots were visualized using the Clarity Western ECL kit from Biorad.

#### Northern blot

Total RNA was extracted from 3 cortices for each sample using Trizol reagent. The aqueous phase was subsequently extracted with phenol-chloroform-isoamyl alcohol (25:24:1; Sigma), followed by an additional chloroform extraction. The RNA was then precipitated using 2-propanol.

Total RNA was separated on a 1.2% agarose gel containing 1.2% formaldehyde in Tri/Tri buffer (30 mM triethanolamine, 30 mM Tricine, pH 7.9). The migrated RNA was transferred overnight to a Hybond N+ nylon membrane by passive transfer. Prehybridization was performed for 1 hour at 45°C in a buffer containing 6× SSC, 5× Denhardt’s solution, 0.5% SDS, and 0.9 g/mL tRNA. Following prehybridization, the membrane was incubated overnight with 5′-radiolabeled oligonucleotide probes. The sequences of the probes used were: ITS1: 5’-GCT-CCT-CCA-CAG-TCT-CCC-GTT-AAT-GAT-C-3’; ITS2: 5’-ACC-CAC-CGC-AGC-GGG-TGA-CGC-GAT-TGA-TCG-3’;18S: 5’-CCG-GCC-GTC-CCT-CTT-AAT-CAT-GGC-3’ and 28S: 5’-CCC-GTT-CCC-TTG-GCT-GTG-GTT-TCG-CTG-GAT-A-3’. After hybridization, the membrane was washed twice for 10 minutes in 2 × SSC, 0.1% SDS, followed by one wash in 1× SSC, 0.1% SDS. The membrane was then exposed to light, and signals were captured using a Typhoon Trio PhosphoImager (GE Healthcare). Signal intensities were quantified with MultiGauge software.

#### En face immunohistochemistry

F-actin immunodetection was performed to delineate cell boundaries. Briefly, mouse embryonic brains were fixed in 4% w/v paraformaldehyde (PFA) (Sigma-Aldrich, France). Cortical explants were dissected and incubated 15 min at RT in PBST 0.1% (PBS 1X containing 0.1% Triton X-100). Explants were then incubated 2 hours (h) at room temperature (RT) in blocking solution (PBS 1X, 0.1% Triton X-100, 10% Normal goat serum). Washes were performed in PBST 0.1%, and then the explants were incubated with Alexa Fluor 633 Phalloidin (1:400, Life Technologies) in PBST 0.1% O/N at 4°C. Extensive washing was performed in PBST 0.1% and PBS 1X before mounting the explants with Fluoromount G (Invitrogen) with the ventricular surface (VS) up to obtain an en face view of the ventricular side of the cortex. Fluorescently stained sections were imaged with a confocal TCS Leica SP5-II microscope. **Analyses:** Confocal images covering the depth of apical endfeet were acquired with 100X magnification and 1024x1024 resolution. A randomly chosen ROI (50 x 50 μm) of a maximum intensity projection image was analyzed per image and three images were used per embryo. Images were analyzed using the ImageJ. The Tissue Analyser plugin was used to create a segmentation trace of each apical image, which was then utilized to measure the area of each apical extremity segment in the ROI. The percentage of segments in a range of lower to higher areas in a defined ROI was then computed.

### In utero electroporation (IUE)

Timed-pregnant mice (E14.5) were anesthetized with isoflurane (4% during induction and 2–2.5% during surgery) and embryos were revealed within the intact uterine wall after sectioning the abdomen. Embryos were constantly hydrated with NaCl 0.9% (B. Braun). A solution containing plasmid DNA (1 µg/μl,) and 20 % wt/vol fast green in sterile endo-free water was injected into the lateral ventricles of the embryos. Forcep electrodes (System CUY650P5 NepaGene Co) were placed around the embryo head at a 45° angle and plasmids were electroporated by discharging a 4,000-μF capacitor charged to 35 V (five electric pulses of 50 ms with 950 ms intervals) with a CUY21 NepaGene electroporator. The embryos were then placed back in the abdominal cavity for 24 h prior to subsequent analyses. Embryonic heads were harvested and fixed overnight with 4% PFA at 4°C.

Brain sections were fluorescently immunolabeled (see below) with antibodies detecting GFP in electroporated progenitors. Images were acquired at room temperature with a TCS Leica SP5-II confocal microscope, with analyses focused on the future somatosensory cortex. 40X and 63X objectives were used and controlled by LAS-AF software for acquisition (Leica).

### Embryonic brain collection and sectioning

Females were sacrificed by cervical dislocation and embryos were collected. Brains were fixed overnight with paraformaldehyde (PFA) 4% and then rinsed and stored with phosphate-buffered saline 1X (PBS). For vibratome sectioning, brains were placed in an inclusion of 10% sucrose and 7.5% agarose in PBS 1X. Brains were cut in 70-μm thick coronal sections using a vibrating blade microtome (Leica VT1000 S). For cryostat sectioning, brains were extracted, washed in PBS, and cryoprotected overnight serially in 15% and 30% sucrose. Brains were embedded in an embedding chamber using cryomedium Neg-50 (Ref 6502; Epredia), frozen under isopentane and dry ice, and cryo-sectioned at 20 µm with a Cryostar NX70 (HOMVPD, Microm).

### IUE-TRAP in aRG

#### IUE and polysome extraction

Embryos were electroporated with pCAG-EGFP-Rpl10a as described above and after 24 h the cortices were microdissected and snap-frozen. Polysome extraction was performed following the protocol described in the Mazaré et al. 2020, except that cortices were homogenized in a 2 ml Teflon-glass homogenizer with 20 strokes instead of 12 for the total brain. GFP-fused aRG polyribosomes were immunoprecipited with anti-GFP antibodies and protein-G-coupled magnetic Dynabeads. At the end of the procedure, immunoprecipitated polyribosomes were eluted by boiling the beads in 20 μL of 0.35 M KCl buffer with 5X Laemmli buffer for 5 min. Ribosome-bound mRNAs were purified using the RNeasy Lipid tissue kit (QIAGEN). Three replicates per condition were prepared, and two embryos at least were pooled per replicate.

#### Library preparation and sequencing

Libraries were prepared and sequenced by iGenSeq, Institut du Cerveau, Paris, France. Briefly, mRNA library preparation was realized following manufacturer’s recommendations (SMART-Seq v4 Ultra Low Input RNA Kit from Takara). Each library prep was controlled on AGILENT tapestation 4200. Final samples pooled library prep were sequenced on ILLUMINA sequencer with 300 cycles cartridge, corresponding to 2x70Millions 150 bases reads per sample after demultiplexing.

#### Bioinformatic analyses

RNA-Seq data analysis was performed by GenoSplice technology (Paris, France). The RNA-seq gene expression data and raw fastq files are available on the GEO repository under the accession number GSE313303. Analysis of sequencing data quality, reads repartition (*e.g.*, for potential ribosomal contamination), inner distance size estimation, genebody coverage, strand-specificity of library were performed using FastQC v0.11.2, Picard-Tools v1.119, Samtools v1.0, and RSeQC v2.3.9. Reads were mapped using STAR v2.7.5a (Dobin et al. 2013) on the mouse mm39 genome assembly and read count was performed using featureCount from SubRead v1.5.0 and the Human FAST DB v2022_1 annotations. Gene expression was estimated as described previously (Paillet et al. 2021). Only genes expressed in at least one of the two compared conditions and covered by enough uniquely mapped reads were further analyzed. Genes were considered as expressed if their FPKM value was greater than FPKM of 98% of the intergenic regions (background). At least 50% of uniquely mapped reads was required. Analysis at the gene level was performed using DESeq2 (Love et al. 2014). Genes were considered differentially expressed for absolute value of log2 fold-change ≥ 1 and adjusted p-values (FDR) ≤ 0.05. Pathway enrichment analyses and GSEA analysis (Subramanian et al. 2005) were performed using WebGestalt v0.4.4 (Liao et al. 2019) merging results from up-regulated and down-regulated genes only, as well as all regulated genes. Pathways and networks were considered significant with p-values ≤ 0.05.

### Puromycin treatment

Pregnant dams received intraperitoneal puromycin injections (50 mg/kg, Puromycin, Sigma P7255). One hour later, developing tissues were obtained and sectioned to a thickness of 70 mm using a vibratome. Puromycin signals were detected with a primary antibody. Images were taken at 40X (Confocal microscope) and fluorescence intensity was quantified using FIJI (ImageJ). For each sample, puromycin intensity along the ventricular zone was measured and averaged.

### Immunohistochemistry

Immunohistochemistry for all other experiments was performed on floating brain slices. These were permeabilized with 1X PBST (0.1% Triton X-100) for 15 min. After washes, blocking was performed for 1 h at room temperature (RT) with 1X PBS containing 10% normal goat serum (NGS) and 0.1% Triton X-100 before incubation O/N at 4°C with the primary antibody. After extensive washes, sections were incubated with the secondary antibodies for 2 h at RT protected from the light. This was followed by 15 min incubation in Hoechst stain (1: 10,000; Thermo Fisher Scientific) prior to washing with 1X PBS. Brain slices were mounted using Fluoromount G (Invitrogen). Images were acquired at room temperature with a TCS Leica SP5-II confocal microscope with analyses focused on the future somatosensory cortex. Fluorochromes are as described above in the antibody section. 40X (NA = 1.25–0.75) and 100 X (NA = 1.44) objectives were used and controlled by LAS-AF software (Leica). Minimum contrast adjustment was performed using ImageJ software. For Fbl labeling, antigen retrieval was performed by incubating the sections in sodium citrate 10 mM pH 6 at 95°C for 20 min and allowing them to cool down before blocking.

### Image acquisitions

As mentioned in the above methods, acquisitions of immunolabeled brain sections and plated cells were carried out using confocal microscopes Leica SP5. For vibratome and cryostat brain imaging a total of z = 10 µm was imaged. For cells, a total of z = 5 µm was imaged. Hoechst (DAPI, 359 nm), Alexa Fluor 488 (S32354; Invitrogen), Alexa Fluor 555 (4413; Cell Signaling), and Alexa Fluor 633 (A21052; Invitrogen) were used.

### Statistical analysis

The sample size selection for experiments was based on both published and previous pilot studies considering the sensitivity of the applied approaches. When possible, data were collected and analyzed in a blind manner to the experimenter. One main experimenter performed each experiment. Statistical tests were carried out using GraphPad Prism 10. Normality and homogeneity of variances were tested using either a D’Agostino-Pearson, Shapiro-Wilk or KS normality tests, or data distribution was assumed to be normal but this was not formally tested, depending on the number of samples. Significance was established with P value ≤ 0.05. For each experiment, the statistical test used (Mann Whitney, T-test or Two-way Anova) is described in the figure legend, as well as the number of individuals analysed. Data were collected and processed randomly.

## Acknowledgements

We thank Ana Uzquiano for her starting data, helping to design the initial project. We are grateful to the members of the Francis lab, Dr. A. Baffet and Dr. J.B. Brault for comments and discussions. We thank L. Broix for help in analyses and interpretation of MitoTracker. We thank P. de la Grange (GenoSplice) for bionformatic analyses. We thank the IFM animal experimentation facility and cellular and tissue imaging platforms, supported also by the *Région Ile de France* (including the DIM CBRAINS project) and the FRC Rotary. We also thank the TAAM (CDTA, Orléans) for aid with animal maintenance. We thank the METi Platform (Dir. S. Balor, member of the national infrastructure France-BioImaging supported by the French National Research Agency ANR-10-INBS-04), and the CALMIP HPC facility (Toulouse, France: Dir JL Estivalezes)

## Author contributions

V.V. conceived, designed, performed, and supervised animal and in vitro experiments, analysed data, and wrote the manuscript. K.C. helped with mice genetic characterization and brain phenotype analyses. C.C.D. and M.S. performed EM experiments. R.T. and M.C.S. helped performing TRAP experiments. S.L. and C.P.C. conceived Ribo Mega-SEC experiments and S.L. helped performing Ribo Mega-SEC and performed WB and NB. R.B. helped perform live imaging experiments. R.S. and C.P.C. conceived electron tomography experiments, which were performed by D.R., V.S. and R.S. C.J, M.M.M. and S.C. conceived ECL-based immunoassays and S.C. helped perform ECL-based immunoassays F.F. initiated the project, conceived, designed, and supervised experiments, helped with data analyses and interpretation, and wrote the manuscript.

